# The infinite alleles model revisited: a Gibbs sampling approach

**DOI:** 10.1101/2021.07.21.452479

**Authors:** Marc Manceau

## Abstract

The SARS-CoV-2 outbreak started in late 2019 in the Hubei province in China and the first viral sequence was made available to the scientific community on early January 2020. From there, viral genomes from all over the world have followed at an outstanding rate, reaching already more than 10^5^ on early May 2020, and more than 10^6^ by early March 2021. Phylodynamics methods have been designed in recent years to process such datasets and infer population dynamics and sampling intensities in the past. However, the unprecedented scale of the SARS-CoV-2 dataset now calls for new methodological developments, relying e.g. on simplifying assumptions of the mutation process.

In this article, I build on the *infinite alleles model* stemming from the field of population genetics to develop a new Bayesian statistical method allowing the joint reconstruction of the outbreak’s effective population sizes and sampling intensities through time. This relies on prior conjugacy properties that prove useful both to develop a Gibbs sampler and to gain intuition on the way different parameters of the model are linked and inferred. I finally illustrate the use of this method on SARS-CoV-2 genomes sequenced during the first wave of the outbreak in four distinct European countries, thus offering a new perspective on the evolution of the sampling intensity through time in these countries from genetic data only.

## 1 Introduction

The concept of *descent with modification* is central in modern biology, where biological entities evolving across various spatial and temporal scales (e.g. cells, individuals, species) can be seen as atomic particles carrying molecular sequences, that are passed on to their descent, while accumulating small gradual changes. As a result, the patterns of genetic differentiation obtained in a sample of particles depend on the underlying population dynamics, and can be analysed to retrieve information on this unobserved population dynamics. This is the aim shared by two related fields called *population genetics* and *phylodynamics*.

### Population genetics and phylodynamics

Molecular sequences are nowadays routinely being collected and analyzed throughout the tree of life, to address a wealth of biological questions, across fields such as ecology, anthropology, macroevolution, developmental biology, or epidemiology. In this manuscript, I focus on methods designed to investigate the *population dynamics* of a system through the analysis of genetic polymorphism. These methods have been applied across plenty of temporal and geographical scales, e.g. in ecology to study the population size trajectory of species (Parag et al. 2021), in epidemiology to estimate the prevalence of an infectious disease from sequences of pathogens sampled during an outbreak (Stadler et al. 2013), or in paleontology to study the species diversity trajectory of a clade over macroevolutionary time-scales (Morlon et al. 2011). While both fields address similar questions and may seem intertwined, their methodologies remained quite distinct, giving rise to two branches in the literature.

Population genetics approaches primarily aim at studying genetic variation within populations through time, based on genetic data. The recognition of the central influence of demography on genetic variation fostered the development of statistical methods aiming at inferring past demography from observed genetic polymorphism. The field has been very active since the beginning of the 70s, and most of the early theory is now digested in textbooks presenting the coalescent, with or without demography complications (Tavaré 2004; Hein et al. 2004; Durrett 2008). This early work relied on simplifying assumptions such as the *infinite alleles* or *infinite sites* models, when genetic data has been sampled at a single point in time. Elegant analytical developments of the probability distribution of summary statistics were derived, allowing one to investigate, e.g., population growth (Kuhner et al. 1998), population structure (Beerli and Felsenstein 1999), selection (McDonald and Kreitman 1991), or the presence of recombination (Hudson 1983). Contemporary empirical applications usually deal with more complicated demography scenarios and samples taken at multiple points in time, and have thus adopted two different strategies. First, some studies rely on Principal Component Analysis or summary statistics that have been previously derived in very simple settings (Novembre et al. 2008). This approach is extremely fast and appropriate for an initial exploration of the dataset, still it lacks a quantitative aspect. Second, the rise of computational power fostered the development of Approximate Bayesian Computation to fit parameter-rich models using computationally intensive procedures (Skoglund et al. 2014; Kim et al. 2017).

Phylodynamics approaches stem from the field of phylogenetics, which aims at reconstructing the ancestral relationships between individuals, together with their evolutionary parameters, based on genetic data. In this field as well, researchers have acknowledged the key role of the demography in shaping the phylogenetic tree and hence the observed molecular patterns. This in turn promoted the rise of a subfield called *phylodynamics*, aiming at inferring the demography using molecular sequences, by integrating over precise phylogenetic relationships. The two main demography frameworks used in the field are (i) the coalescent, borrowed from Kingman (1982)‘s work in population genetics and (ii) birth-death processes, relying on seminal results by Kendall (1948) and Nee et al. (1994). Compared to population genetics methods, there has been a cultural change towards more precise estimation relying on computationally intensive Bayesian inference methods. These methods rely on many superimposed model layers, among which e.g. models of clock evolution (Lepage et al. 2007), models of across-locus variation (Lartillot and Philippe 2004), and models of molecular substitution (Lanave et al. 1984). Moreover, phylodynamic methods have been developed to take into account serially sampled molecular data (Stadler 2010). Population dynamics has been modeled either in a coalescent framework using e.g. time-varying population size (Pybus and Harvey 2000; Drummond et al. 2002; Pybus et al. 2003), or in a birth-death framework, where the population is already free to fluctuate with constant birth and death rates, but larger variations can be allowed using time-varying parameters (Morlon et al. 2011; Stadler et al. 2013). As an alternative to time-dependent processes, some studies have attempted to introduce diversity-dependence processes in either a coalescent framework (Volz et al. 2009), or a birth-death framework (Etienne et al. 2012; Leventhal et al. 2013). Population structure can be modeled in a coalescent framework with discrete demes exchanging genes through migration (Ewing et al. 2004; Vaughan et al. 2014; Müller et al. 2017). In a birth-death process, structure is modeled using so-called multi-type birth-death processes, where different types are associated with different birth and death parameters, and individuals from a given type can either give birth to other types or directly change type (Maddison et al. 2007; Beaulieu and O’Meara 2016; Maliet et al. 2019; Barido-Sottani et al. 2020). Finally, methods have been developed to jointly consider occurrence and molecular data. In a coalescent framework, occurrences are assumed to be the result of a Poisson sampling process among the total population (Rasmussen et al. 2011; Parag et al. 2020). In a birth-death process, an individual can be sampled and *sequenced* at a given rate -in which case it appears in the tree -or sampled without being sequenced at another rate -in which case it is a simple occurrence (Vaughan et al. 2019; Gupta et al. 2020; Manceau et al. 2021).

### Motivating example

In this paper, I focus on inferring population dynamics for biological systems where (i) genetic polymorphism is sampled through time, and (ii) an ever increasing amount of sequences are being collected, challenging state-of-the-art methods for phylodynamic analysis.

The current SARS-CoV-2 pandemic provides the archetypal dataset that I propose to model. The outbreak survey started in late 2019, and the first viral genome was already published and made available for research on the 10th of January 2020. New sequences followed at an outstanding rate. By early May 2020, already more than 10^5^ viral sequences were available from across the world. A bit less than a year after, 10^6^ sequences have been reached before early March 2021. Developing statistical tools capable of keeping up with the pace of data acquisition thus represents a methodological challenge.

SARS-CoV-2 genomes have already been used to address a number of epidemiology-related questions, among which assessing the number and origins of introductions in a given locality (Gonzalez-Reiche et al. 2020; Lemey et al. 2020), the magnitude of super-spreading events (Li et al. 2020), or estimating the reproductive numbers of local outbreaks (Vaughan et al. 2020). Yet, phylogenetics/phylodynamics approaches do not scale well to large numbers of sequences and empirical applications typically require subsampling the original datasets.

The virus genome is approximately 3 × 10^4^ nucleotides long, and its mutation rate, quite heterogeneous across the genome, has been estimated around 22 mutations per year per genome (Hadfield et al. 2018). As a result, new alleles and polymorphic sites of the genome have accumulated in the data at a slow pace. Together with the outstanding number of sequences, this rather slow mutation rate advocates for the use of simplifying assumptions of the mutation process.

### The infinite alleles model

Phylodynamic analyses generally assume a very realistic mutation process. Sequences have a finite number of sites, and each mutation hits a randomly chosen nucleotide, with a realistic substitution process ranging from the Jukes-Cantor to the Generalized Time Reversible model. Selection might even be modelled and nucleotides might have different mutation rates along the sequence. While these realistic models are very well designed to study fine-grain processes or processes happening over long timescales, they do not appear to be the best option to process large numbers of similar sequences. In this manuscript, I take a step back and aim at bringing back into fashion a simplifying assumption that has been traditionally considered in the early days of the neighbouring field of population genetics, namely the *infinite alleles model*. Under this model, each mutation hitting a sequence always creates a new *allele* never observed before. If we imagine that each sequence is a ball and an allele is a colour, genetic data thus simplifies as a sampling record of coloured balls through time, as illustrated in Figure 1 (Durrett 2008).

**Figure 1:**
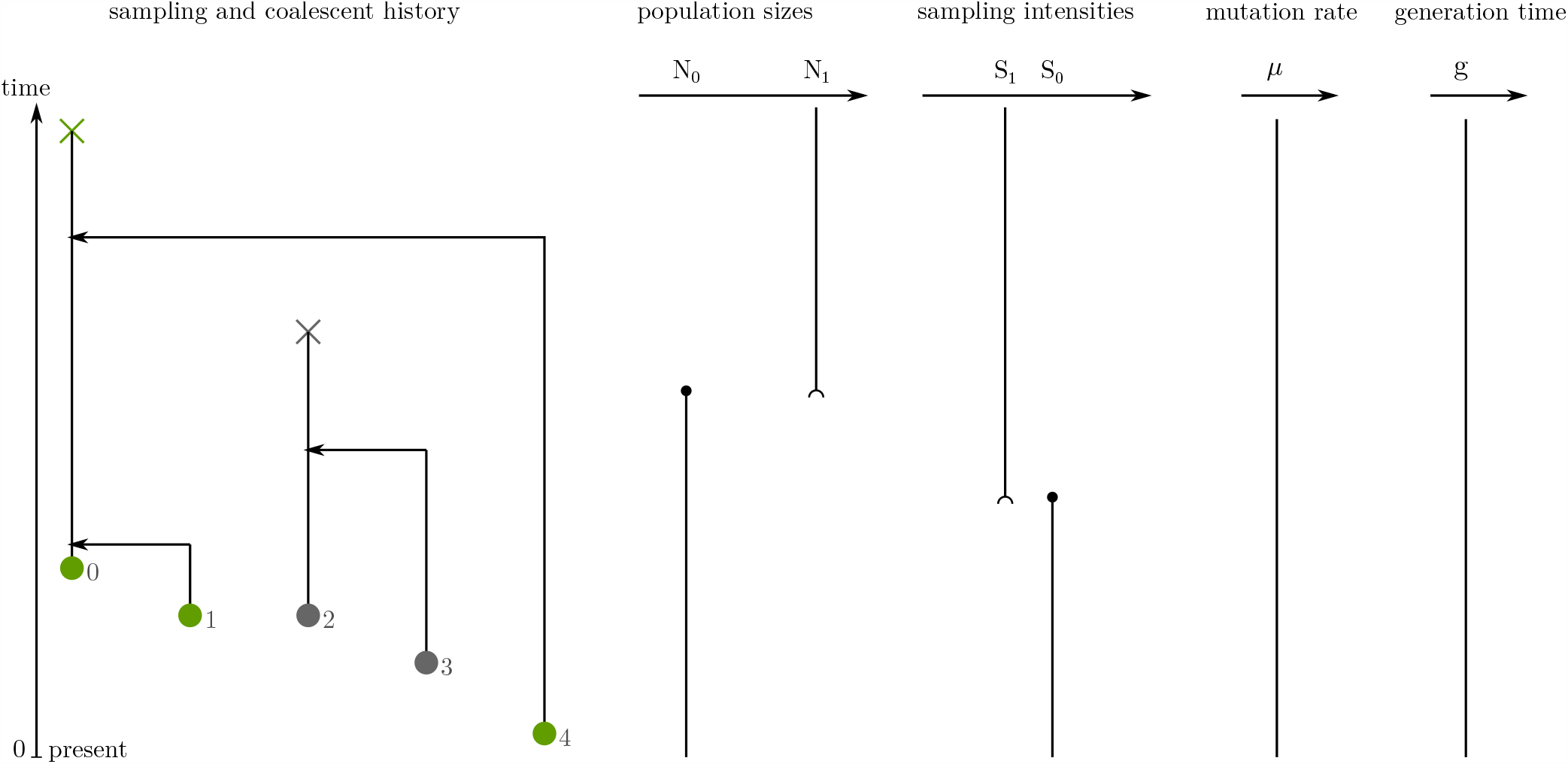
Model notation. Time is oriented from the present towards the past. To the left, individuals, represented as circles at their sampling time, arc numbered in decreasing sampling times order and arc colored according to the allele they belong to. Past coalescent events arc represented with arrows, differentiation events arc represented with crosses. The past coalescent history leads to the allele partition *𝒜 =* {{0,1,4}, {2,3}}. To the right, the parameters of the model that we arc interested in inferring: the piecewise-constant past effective population sizes *N*, the piecewise-constant past sampling intensities S. the constant mutation rate g, and the constant generation time *g*.

Analytical tractability is the main reason why the infinite alleles model is used nowadays. Following the past history of one genetic sequence backward in time, it can either (i) coalesce with another lineage that belongs to the same allele; or (ii) if it is the only representative of its allele, it can find the mutation that gave rise to it. Once this original mutation is found, everything else in the past is forgotten. The infinite alleles model was studied extensively during the golden age of population genetics, in combination with the coalescent model and for sequences sampled at a unique point in time. A closed-form analytical characterization of the probability distribution of the allele frequency spectrum in this setting exists, called *Ewen’s sampling formula* (Ewens 1972).

The dynamics of the colour assemblage through time is informative of the underlying population dynamics that we are interested in inferring. I propose to work under a Bayesian framework, and to rely on population dynamics and sampling process assumptions similar to what has been recently used in phylodynamics. To ensure fast convergence of the Markov Chain Monte Carlo (MCMC) method that is used for inference, the model is (i) carefully built such that data augmentation can be performed efficiently, and (ii) relies on prior conjugacy properties and Gibbs sampling moves. This approach has been successfully applied to other phylogenetics methods before, and has shown a much faster convergence of MCMC methods relying on Gibbs sampling moves as compared to Metropolis-Hastings moves (Lartillot 2006). Further, prior conjugacy properties allow one to build a better intuition on the interactions between different parameters, which proves particularly convenient for the choice of prior distributions. As compared to state of the art phylodynamics methods, I aim at integrating over the unknown ancestral relationships more efficiently, with the hope to warrant dataset analysis on a larger scale.

### Manuscript outline

In Section 2, I introduce in more details the model assumptions, before turning to the inference strategy in Section 3. I then present in Section 4 some sanity checks and validate the inference method on simulated data. An empirical application in Section 5 illustrates the use of the method on the SARS-CoV-2 sequences sampled during the first wave hitting Europe in 2020. Finally, I discuss in Section 6 the results of this paper as well as the future research challenges it opens. This manuscript is released along with the code implementing the method, and details on the implementation and use of the code are provided in Supp. Mat. C.

## 2 Model and notation

I build here on work by Parag et al. (2021) and Karcher et al. (2020), who both consider a sampling process on top of a coalescent model with piecewise-constant effective population sizes. The coalescent process is very conveniently described backward in time, and time will thus be, throughout the manuscript, the calendar time before present, in units of days for empirical applications, with *t* = 0 at present and *t* →∞ in the past.

### 2.1 Model parameters

The model is built around the following four key parameters.

First, the past effective population size is piecewise-constant on a partition of (0, +∞) into *p* successive disjoint intervals 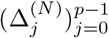 delimited by 0, ∞, and the *p* − 1 times 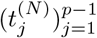, i.e.

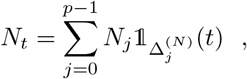

where 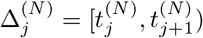, with convention 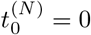 and 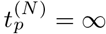.

Second, following Parag et al. (2021), the past sampling intensity is also a piecewise-constant function, on a possibly different partition 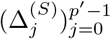 of (0, ∞), delimited by 0, ∞, and the *p*′ – 1 times 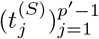, i.e.

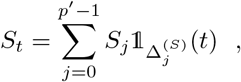

where 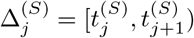, with convention 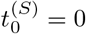 and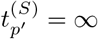.

Last, the mutation rate *μ* and generation time *g* are constant through time.

In a Bayesian framework, these parameters are random variables which are assigned prior distributions. The effective population sizes 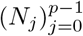 are assumed to be *a priori* distributed according to a Generalized Inverse Gaussian distribution, while the sampling intensities 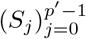 are assumed to be *a priori* distributed according to a Gamma distribution. Finally, the mutation rate *μ* and generation time *g* are respectively *a priori* distributed according to a Gamma and Inverse-Gamma distribution. The choice of these prior distributions will be explained in Section 3, when discussing the posterior inference of these variables.

### 2.2 Past sampling and coalescent history

We assume that the sampling history is given by a Poisson Point Process (PPP) with rate

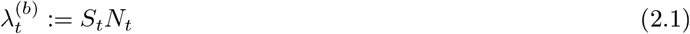

generating the set of ordered sampling times 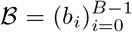 of all individuals. The total number of sampling events is denoted *B*, and individuals are numbered from 0 to *B —* 1 in reverse birth time order. We will also call these sampling times the *birth times* of lineages when considering the history backward in time (hence the name *ℬ*). Lineages begin their backward-in-time journey as singletons {*i*} where *i* corresponds to the individual’s number.

The past history of these lineages is further assumed to follow a standard coalescent with effective population size *N*_*t*_, generation time *g* and differentiation under an infinite alleles model with mutation rate *μ*. That is, while there are *k*_*t*_ lineages alive in the process, the next coalescent (resp. differentiation) event happens with rate,

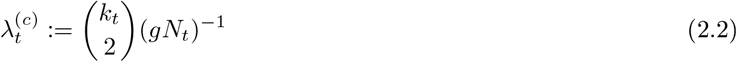

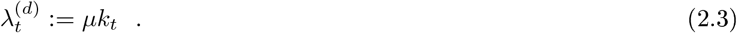

When there is a coalescent event, two lineages *L*_*i*_ and *L*_*j*_ uniformly sampled among the *k*_*t*_ living lineages at that time, are merged together in a unique lineage *L*_*i*_ U *L*_*j*_ When there is a differentiation event, one of the *k*_*t*_ living lineages is uniformly chosen to be killed. Forward in time, a coalescence corresponds to an individual giving birth to another individual, whereas a differentiation event corresponds to the acquisition of an original mutation responsible for the creation of an allele.

The past coalescent history thus generates in particular a partition of individuals into an *allele partition 𝒜* corresponding to the collection of all lineages killed by a mutation. It also generates the times at which differentiation and coalescent events, -jointly referred to as *death* events -happened in history. In order to record these, we take the following approach. Lineages are initially numbered with the same number as the individual’s number it carries. Each coalescence involving two lineages numbered *j < i* at time *t* is considered to kill lineage *i*, and to keep living in lineage *j* (see mows on Fig. 1). By a slight abuse of Irnguage, we call such an event the death of *individual i*, and the time at which a mutation is found is the death time of the very first individual of the allele (see crosses on Fig. 1).

The coalescent history of all individuals is recorded in 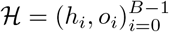, where *h*_*i*_ is the death time of individual *i* and o_*i*_ ∈ {0,1,…, *i*} is the output of the death event, i.e. the number *j* < *i* of the lineage in which lineage *i* is merged if there is a coalescence, or o_*i*_ = *i* if there is a mutation. The total number of alleles is denoted 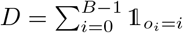. Finally, the record of the *B* birth times *(b*_*i*_) and death times (*h*_*i*_), together with boundaries *t* = 0 and *t* = ∞, yield a partition of the timeline into 2*B* + 1 successive intervals that are denoted 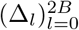. On any such interval Δ_*l*_, the number of lineages remain constant and is denoted *k*_*l*_.

### 2.3 Density of the full history

Figure 1 summarizes all notation introduced so far. Note already that, knowing *ℬ* and *ℋ*, i.e. the full sampling and coalescent history, is enough to know the partition *𝒜* of individuals into alleles. The density of this full past history *ℬ, ℋ* given *N, S, μ, g* is further given by

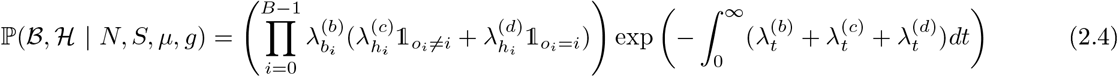

The above density belongs to the exponential family, and thus lends itself well to inference via a Gibbs sampling strategy, with priors that belong to other exponential family distributions. In the next Section, I turn to the description of this inference method.

## 3 Inference method

### 3.1 Observations and inference strategy

Data consist in the observation of the sampling times *ℬ*, together with the partition *𝒜* of the set of *B* individuals into *D* alleles. I aim at inferring the posterior distribution of *N, S, μ, g*, which consists, in a Bayesian framework, in sampling from,

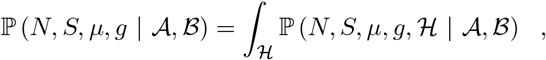

where the right-hand side is augmented using the past coalescent history.

I take a Gibbs sampling approach to design a MCMC which converges to the stationary distribution of the augmented target distribution ℙ *(N, S, μ,g*, ℋ | *𝒜, ℬ)*. To do so, I derive efficient ways to alternatively sample from the following conditional laws,

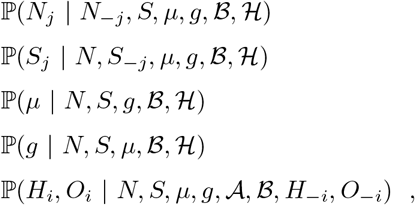

where *N*_−*j*_, *S*_−*j*_ .denote the effective population sizes and sampling intensities over all intervals other than the one numbered *j*, and *(H*_*-i*_, *O*_*-i*_) denotes the death information of all individuals other than the one numbered *i*. Remark here that, on the first three lines, *𝒜* disappeared from the conditioning because *𝒜, ℬ, ℋ* = *ℬ, ℋ*, i.e. knowing the sampling times and the coalescent history is enough to know the allele partition.

Designing an efficient Gibbs sampler in this context relies on the optimization of two critical steps: (i) one needs to derive the posterior distribution of the parameters conditioned on the augmented data; and (ii) one needs to efficiently perform a data augmentation step, i.e. simulate the past history *ℋ* conditioned on the observed data and parameter values.

### 3.2 Prior conjugacy properties for parameters

#### 3.2.1 Effective population size

Recall that *N*_*j*_ is *a priori* independ ent on *N*_*−j*_, *S, μ, g* and is distributed according to a Generalized Inverse Gaussian distribution denoted *𝒢ℐ𝒢(λ, ρ, ψ*). The following will justify this choice of prior. The GIG distribution belongs to the exponential family and is characterized by its density, usually parameterized as,

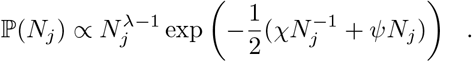

Its posterior is thus given by,

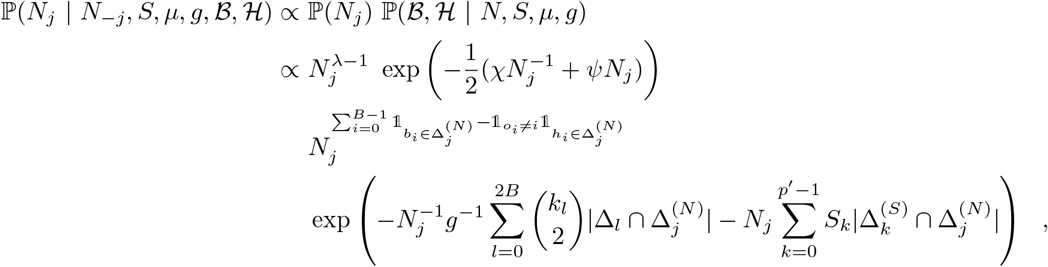

where the last line is obtained by substituting the density of *ℬ, ℋ* using Equation (2.4), before dropping out all terms which do not depend on *N*_*j*_.

This proves that the prior and posterior are conjugate distributions, with

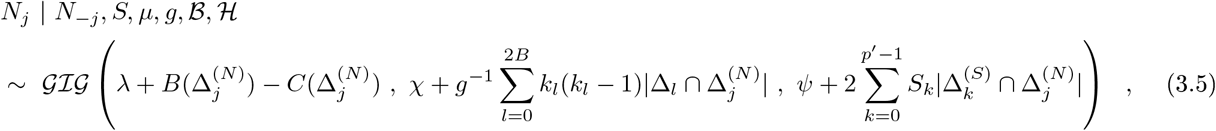

where 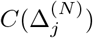 and 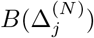 are respectively the number of coalescent and sampling events happening over interval 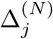.

The choice of a conjugate prior will help (i) simplify the Gibbs sampling process, and (ii) provide a better intuitive understanding of the factors that influence the distribution of *N*.

#### 3.2.2 Sampling intensity

Recall that *S*_*j*_ is *a priori* independ ent on *N, S*_−*j*_, *μ, g* and that S_*j*_ *∼* Γ (*α, β)*. Its posterior is thus given by,

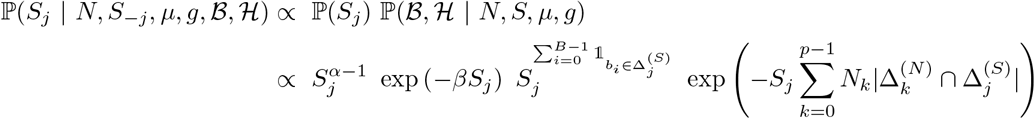

where the last line is obtained again by substituting the density of *ℬ, ℋ* using Equation (2.4), before dropping out all terms which do not depend on *S*_*j*_.

This shows that the prior and posterior are conjugate distributions, with

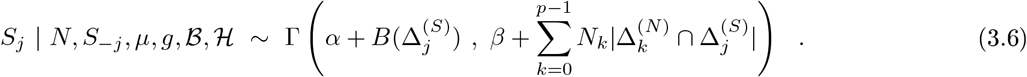

#### 3.2.3 Mutation rate

Recall that, *a priori, μ ∼* Γ (*α, β*). Following the same strategy as above, its posterior is given by,

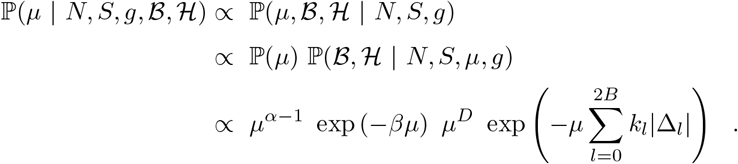

We conclude that the prior and posterior of *μ* are conjugate distributions, with,

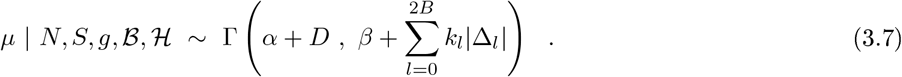

#### 3.2.4 Generation time

Last, recall that *g* is *a priori* distributed according to an Inverse-Gamma distribution denoted Γ ^−1^(*α, β*). Its posterior is given by,

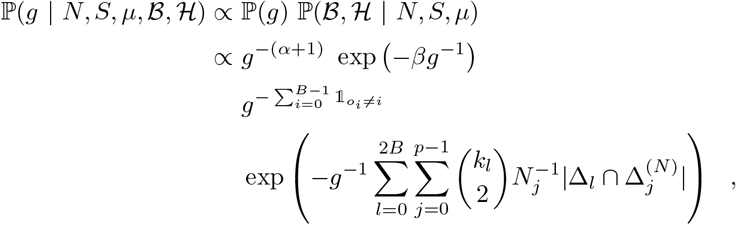

which means that the posterior is

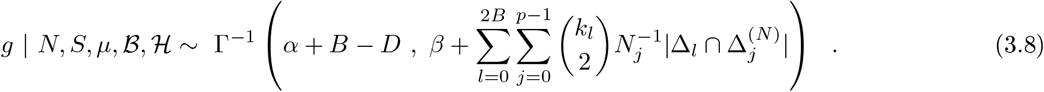

This ends the description of the four conjugate priors used for *N, S, μ, g*. They will be used in the final Gibbs sampler to quickly update the posterior of these variables of interest, provided *ℬ, ℋ* are observed.

### 3.3 Data augmentation with the past coalescent history

Under the assumption of an infinite alleles model with constant effective population size *N* and constant mutation rate *μ*, the distribution of the past history of a sample of n genes taken at a single point in time can be efficiently sampled using the equivalence with the *Hoppe’s urn process* (Durrett 2008). Yet, when sequences are *heterochronous*-i.e. have been sampled through time instead of at a single point in time -and when the population size is not constant anymore, an alternative strategy is needed.

Recall that *H*_*i*_ ∈ ℝ^+^ is the death time of a focal individual *i*, and that *O*_*i*_ ∈ {0,1,…,*i*} is the death output. Death can happen either through coalescence with other lineages (*O*_*i*_ = *j < i*), or through mutation (*O*_*i*_ = *i*). We aim here at sampling death information (*H*_*i*_, *O*_*i*_) of each sequence in turn, conditioned on everything else, i.e. when *N, S, μ, g, 𝒜, ℬ, ℋ*_*-i*_, *O*_*-i*_ are fixed.

In a standard coalescent process, one can draw (*H*_*i*_, *O*_*i*_) from left to right: if a focal lineage *i* sees *k*_*t*_ other lineages already drawn to the left, at time t, it coalesces with one of them at rate *k*_*t*_(*N*_*t*_*g*)^−1^, or finds a mutation at rate *μ*. Yet, when imposing our conditioning, only a fraction of these events leads to the known allele partition *𝒜* and is compatible with the known coalescent history (H_−*i*_,*O*_−*i*_). In particular, the allele partition and past coalescent history excluding i reveal that (i) lineage *i* dies through a coalescent event if and only if it can coalesce with an other lineage to the left, and (ii) lineage i cannot die before being sampled, nor before reaching the very-last coalescence of other lineages with itself, i.e. *H*_*i*_ ≥ *m*_*i*_ := max ({b_*i*_} U {*H*_*j*_, *j* > i such that *O*_*j*_ *= i*}).

A first strategy to draw (*H*_*i*_, *O*_*i*_) conditioned on everything else would thus consist in drawing (*H*_*i*_, *O*_*i*_) without conditioning, while subsequently rejecting simulations which outcome is incompatible with *𝒜*, 𝒪 _−*i*_, *ℋ* _-*i*_. I take here the following alternative approach to avoid rejecting too many simulations. On all intervals Δ_*l*_ = (*t*_*l*_,*t*_*l+*1_) where the total number of lineages *k*_*l*_, remains constant and where *t*_*l*_ ≥ *m*_*i*,_ the probability that *H*_*i*_ falls within the interval conditioned on everything else is computed,

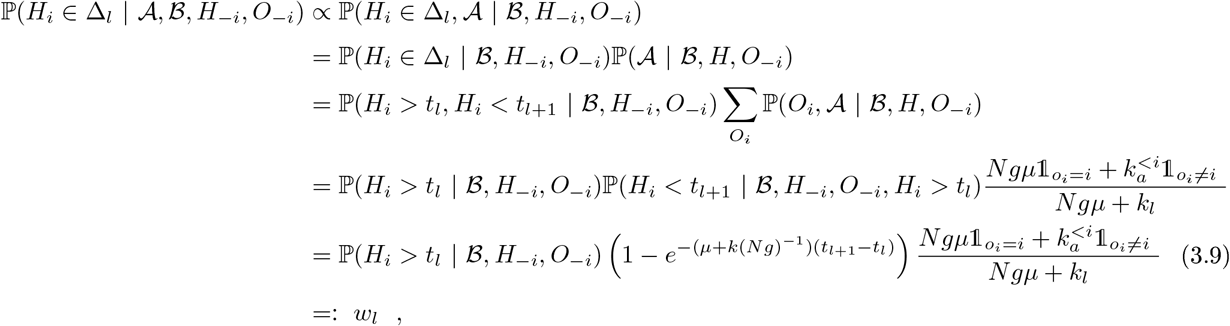

where 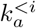 denotes the number of living lineages to the left of *i* belonging to the same allele *a* as *i*.

The above formula is used to recursively compute from bottom to top the weights *w*, associated to all intervals A, above *m*_*i*_. Once these have been computed, we have access to, and can sample from, 𝕡 (*H*_*i*_ ∈ Δ_*l*_, | *𝒜, ℬ, H*_−*i*_, *O*_−*i*_) = *w*_l_/Σ _*k*_ *w*_*k*_. Note in addition that, if lineage *i* satisfies *o*_*i*_ ≠ *i*, then only intervals from time *m*_*i*_ up to time *M*_*i*_ := max *H*_−*i*_ matter, since it must coalesce before the mutation is found. If lineage *i* dies through a mutation, i. e. *o*_*i*_ = *i*, then intervals from time *m*_*i*_ *=* max *H-*_*i*_ up to ∞ must be weighted, since the lineage cannot die before everybody else has coalesced into itself.

Once the interval on which the death event happens has been drawn, it remains to draw from 𝕡 (*H*_*i*_ ∈ *dt* | *H*_*i*_ ∈ Δ_l_,), which corresponds to exponential distribution with rate (*μ*, + *k*_l_(Ng)^-1^) conditioned on happening on Δ_*l*_,. Further, *O*_*i*_ = *i* if the lineage must die through a mutation, or *O*_*i*_ is a uniformly chosen lineage *j < i*, in the same allele as lineage *i*, living on intorvai Δ_*l*_,. The procedure for drawing *(H*_*i*_,*O*_*i*_) is summarized in Algorithm 1 and illustrated in Figure 2.

#### Algorithm 1

Drawing a realization of ℙ (*H*_*i*_, *O*_*i*_ | 𝒜,ℬ,*H*_−*i*_, *O*_−*i*_, *N, S, μ, g*)

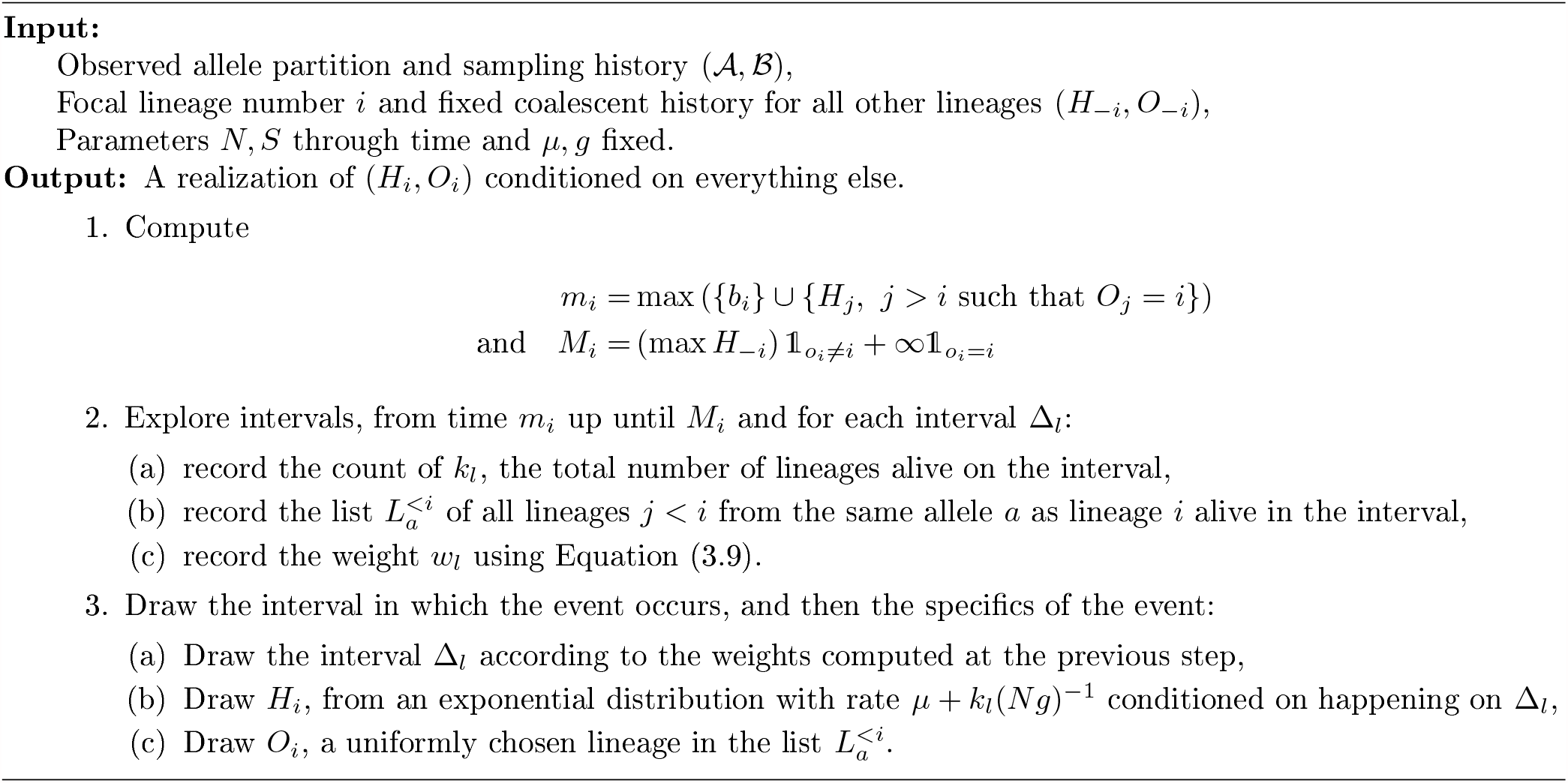

**Figure 2:**
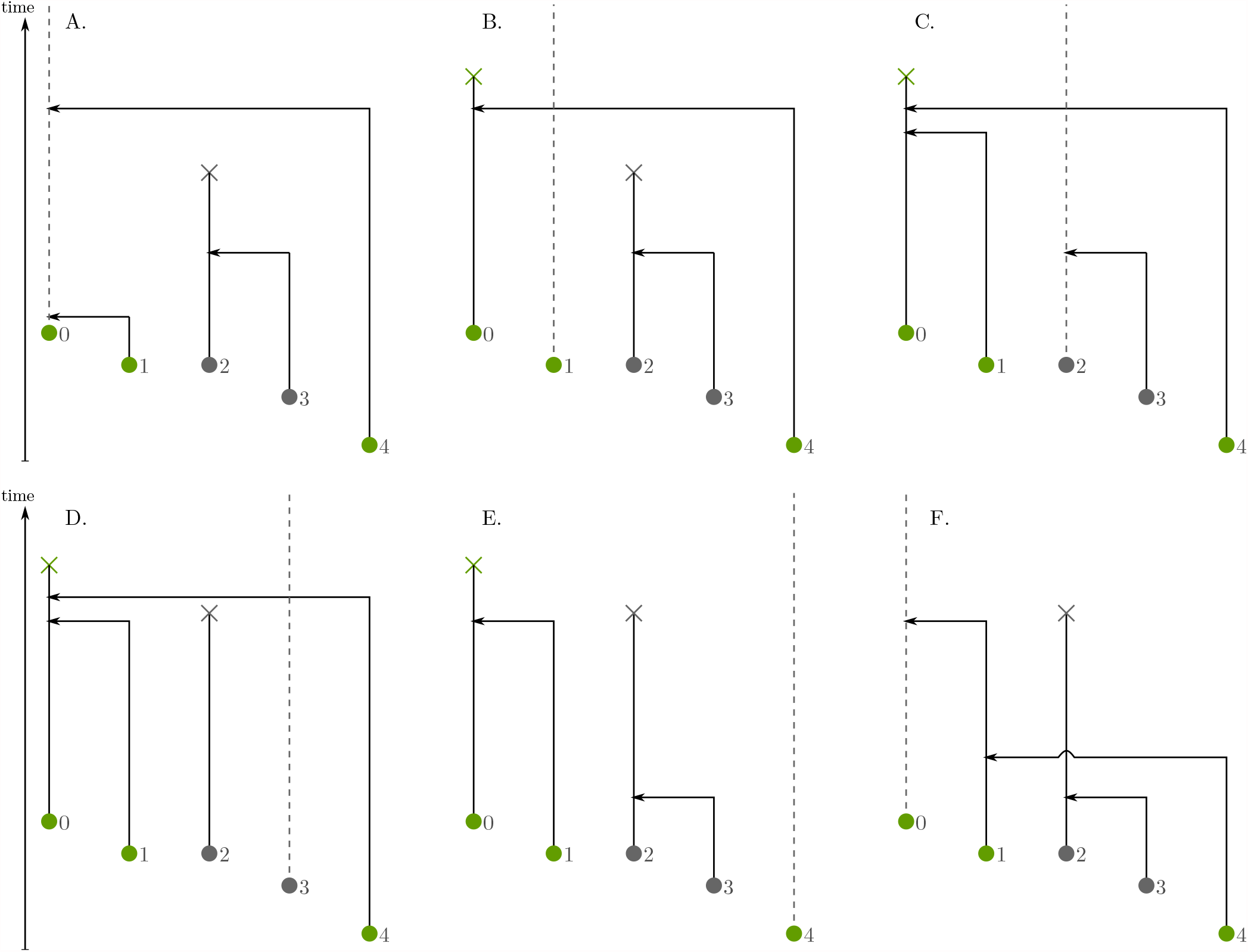
Gibbs sampling approach for simulating the past history of a sample given the allele partition. A-F, Each leaf is considered in turn, and its time of extinction is sampled given the rest of the coalescent history is fixed.

### 3.4 Summary of the Gibbs sampler

The final Gibbs sampler is a simple MCMC that is initialized using the priors of *N, S, μ, g*, before relying on repeated updates of the four variables *N, S, μ, g, ℋ*, using their conditional probabilities. Algorithm 2 summarizes these steps.

#### Algorithm 2

Gibbs sampling of the target distribution ℙ (*N, S, μ, g*, | 𝒜,ℬ)

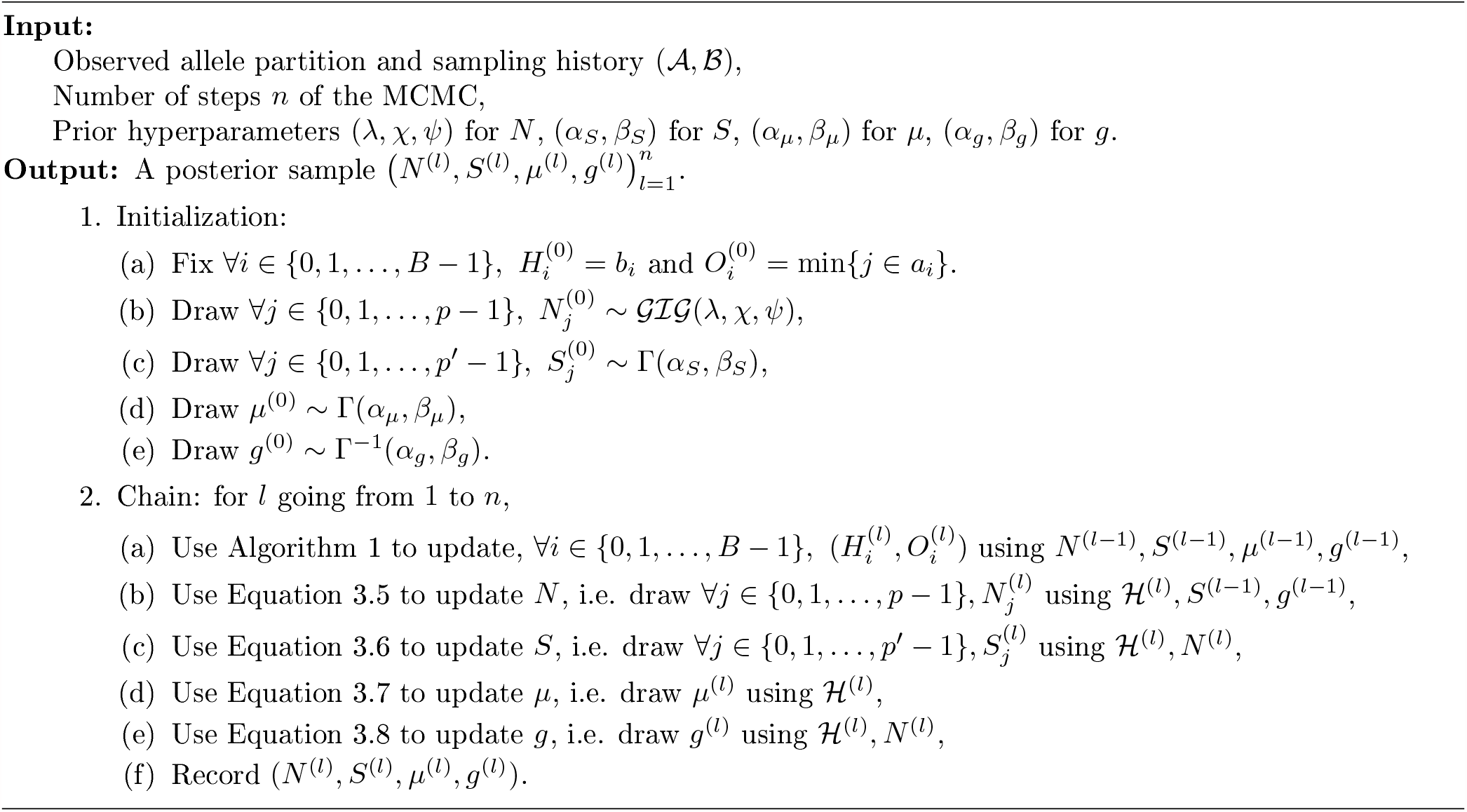

In the next Section, I validate the behaviour of this MCMC on simulated datasets before applying it to empirical datasets in Section 5.

## 4 Numerical validation of the method

### 4.1 Past coalescent history

I first aim to validate the procedure described in Section 3.3 and the implementation of Algorithm 1. To do so, I wrapped Algorithm 1 in a minimalist Gibbs sampler aiming at sampling from 𝕡 (*ℋ \ 𝒜, ℬ, N, S, μ, g*). I fix *ℬ, 𝒜* as well as all parameters of the model N, S, μ, g. I then use a simplified version of the MCMC described as Algorithm 2, where all updates concerning *N, S, μ, g* are skipped. To draw one observation of H, I performed *n* = 50 complete cycles of leaf updates, and kept only the last state reached by the MCMC.

I compare the distribution obtained using this procedure with the distribution of *ℋ* obtained by a naive rejection algorithm, consisting in simulating the coalescent process backwards in time, while rejecting outcomes that do not satisfy *𝒜*. The distributions are compared based on summary statistics computed on 10^4^ samples drawn in both distributions: (i) the proportion of samples having a mutation in successive death events, as well as (ii) the distribution of death times. Note that the dataset must be small, for the rejection algorithm very quickly becomes computationally too intensive to be used. Figure 3 illustrates the perfect agreement between both distributions, on a toy dataset with *N* =10 *g* = 0.1, *μ =* 1.5, and two alleles respectively joining individuals sampled at times (0; 0.2; 0.5) and (0.3,0.7).

**Figure 3:**
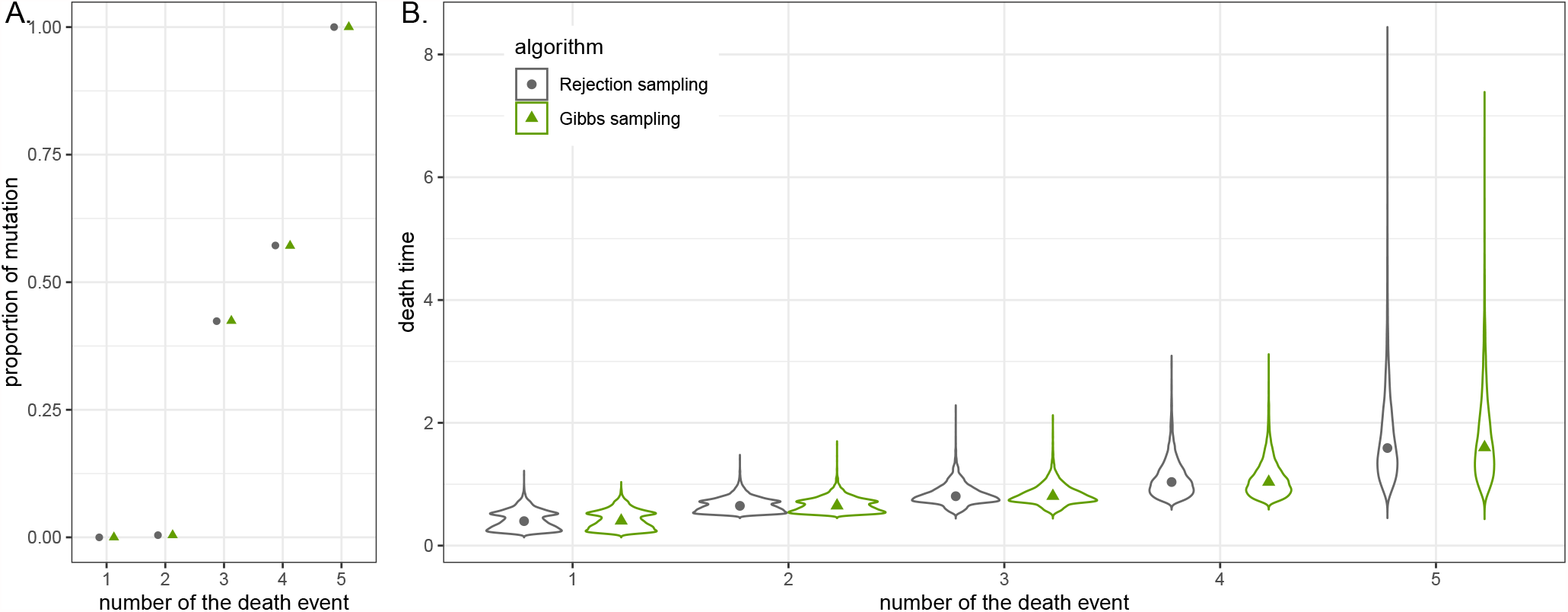
Comparison of the distribution of *ℋ* conditioned on *𝒜* obtained using a naive rejection algorithm or a custom Gibbs sampler relying on Algorithm 1. The number of the death event in x-axis refers to an order by increasing death time. A. Proportions of mutations in each death event. B. Distributions of death times.

### 4.2 Simulation-based calibration

The MCMC implementation of Algorithm 2 is further validated against simulated data using the Simulation-Based Calibration (SBC) method described by Talts et al. (2018) and summarized hereafter.

First, priors arc fixed as follows, with hyperparameter values ensuring that the number of samples remains relatively small. The effective population size is piecewise-constant over *p* = 2 intervals, with *N*_*0*_*;N*_*1*_ *∼ 𝒢 ℐ𝒢 (λ = 4; χ* = 0; *ψ* = 0.08) respectively over intervals (0; 20) and (20; +∞). The sampling intensity is piecewise-constant over *p*′ = 3 intervals, with *S*_0_; *S*_1_ having a Γ (4,1000) prior respectively over intervals (0; 10) and (10; 40) and *S*_3_ = 0 on (40; +∞). The mutation rate and generation time are respectively assigned a prior *μ ∼* Γ (4; 400) and *g* ∼ Γ^−1^(10; 10).

Second, 10^4^ parameter sets are sampled from these distributions and for each parameter set, the sampling history ℬ well as the allele partition *𝒜* are sampled according to the model.

Third, for each simulated dataset, the posterior distribution of *N*; *S*; *μ*; g conditioned on *𝒜*; *ℬ* is sampled using the Gibbs sampler described as Algorithm 2, while using the same priors that were used for the simulation. I ran the MCMC for a total of 10^4^ steps, discarded a burn-in of 10^3^ steps at the beginning and recorded one state every 100 steps over the remaining steps.

Finally, Figure 4A shows the proportion of datasets *p*_*α*_ for which a credible interval with level *α* of the posterior distribution contains the true simulated data, for 9 values of *α* evenly spaced on (0; 1). The good match between *p*_*α*_ and *α* indicates that the MCMC correctly samples the posterior distribution. This is further confirmed in Figure 4B, showing the histogram of the rank statistic associated with the 10^4^ experiments. Here, the rank statistic associated to one experiment refers to the number of samples from the posterior being less than the true value. Under the null hypothesis that we are sampling the true posterior distribution, the histogram of rank statistics should be uniform, as illustrated in Figure 4B (Talts et al. 2018).

**Figure 4:**
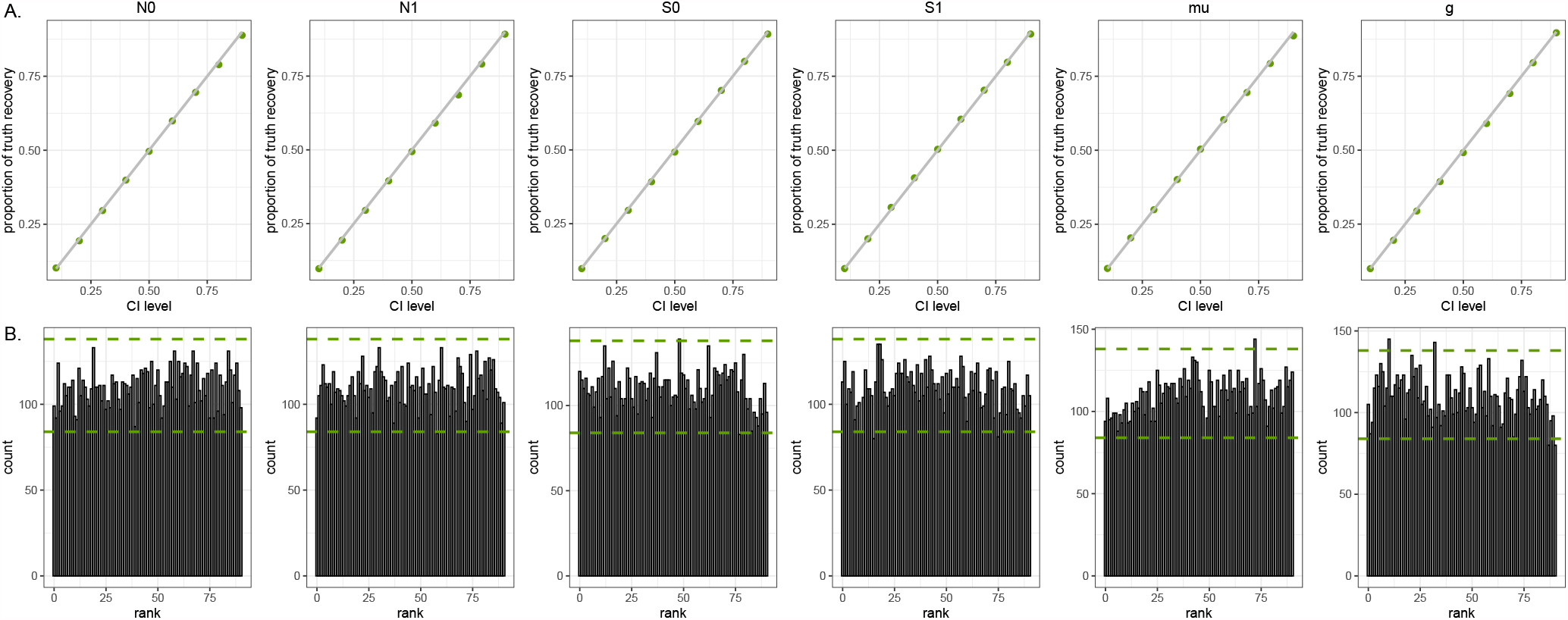
Results of the SBC analysis described in the main text, validating the Gibbs sampler. A. Proportion of datasets which 100*α*% posterior credible interval recovers the true parameter, for different values of *α*. B. Histogram of rank statistics. Horizontal dashed lines indicate the 99% CI of the bar heights if the histogram is uniform.

### 4.3 Running time assessment

I now turn to an assessment of the running time of the method on realistic datasets. I simulated datasets using a timeline cut into *p = p*′ = 5 intervals with varying *N* and *S* values, while fixing the same hyperparameters on all intervals, so as to get datasets with total sample size *B* regularly spaced on a log scale between 10 and 2500. On each of these, I ran 10^4^ steps of the Gibbs sampler, discarded the first 10^3^ steps, and recorded the running time and posterior samples.

Let us focus first on the running time of each of the different updates. Since for each of *B* sequences, the update of the death event requires to compute the weights associated to a number of intervals of the order of B, the update of the coalescent history is expected to scale in *O*(*B*^2^) and to be the bottleneck of the MCMC. Figure 5A presents estimates of the running time depending on B, using the code released along this article on a laptop. It confirms that the update of the coalescent history -and so, a step in the MCMC -scales in *O*(*B*_2_).

**Figure 5:**
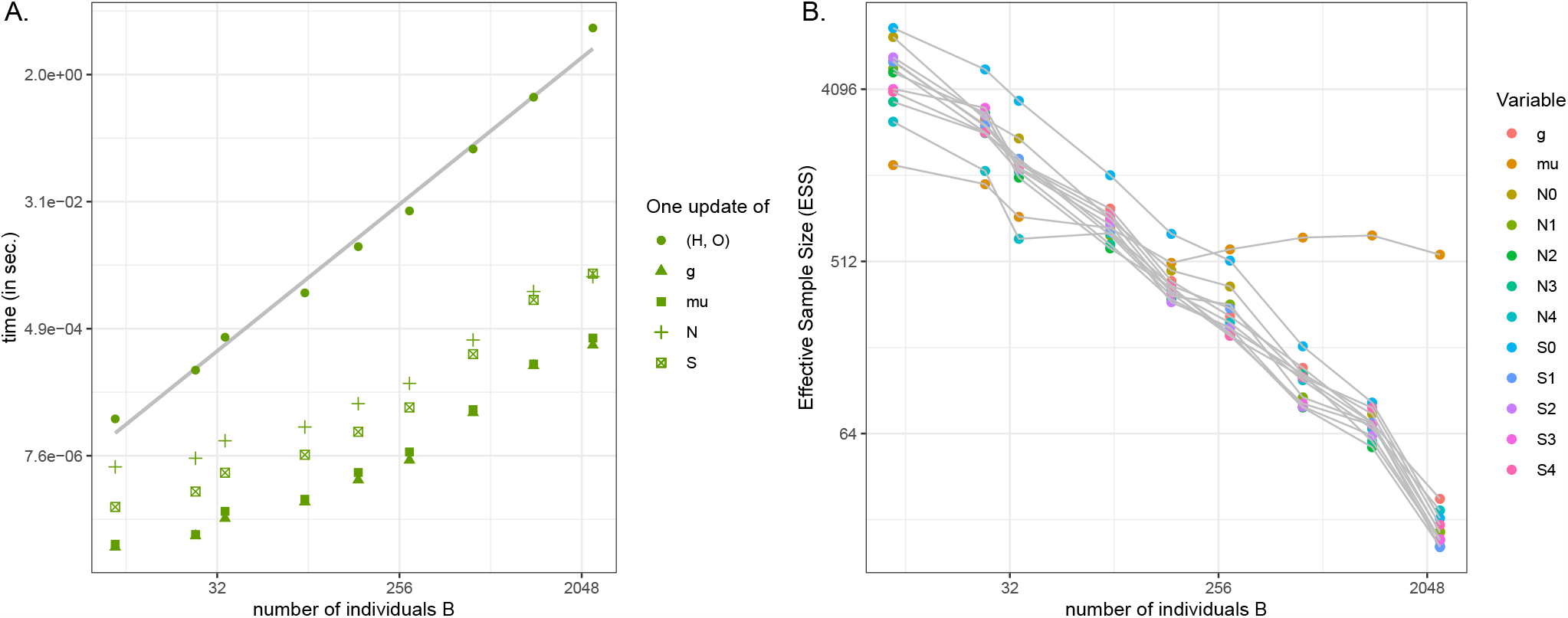
Estimating the running time of the method on simulated datasets. A. The running time of one update of each of the four variables shows that the update of the coalescent history is the bottleneck, that scales in *O*(*B*^2^). B. The ESS of *μ, g* and each of the piecewise-constant values of *S* and *N* as a function of the dataset size suggests that reasonable (i.e. > 100) ESS values for datasets up to *B ∼* 10^3^ can be obtained using ∼ 10^4^ MCMC steps.

Running a MCMC moreover requires to perform these updates repeatedly during a certain number of steps, in order to (i) escape the burn-in phase of the MCMC; and (ii) collect enough samples from the posterior to characterize it. When the samples collected through time are highly correlated, it is said that the chain is *mixing poorly*, and more steps are typically required. In order to provide a rough idea of the expected mixing behaviour on simulated datasets, I computed estimates of the effective sample size of 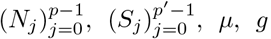. The results are shown in Figure 5B. Combined with the running time assessment, it conveys the idea that the current implementation, without further approximation or numerical optimization will likely not be useful to process more than ∼ 10^4^ sequences.

Having in mind the rough behaviour of the method, I illustrate its use in the next-Section on empirical datasets encompassing ∼ 10^3^ sequences.

## 5 Empirical application

### 5.1 Data collection and preprocessing

Sequences collected in Switzerland (CH), Germany (DE), France (FR) and Italy (IT) between December 1st 2019 and June 1st 2020 have been downloaded on the GISAID website on the 7th of June 2021, while requiring the three options of (i) *complete* sequences, (ii) *high coverage* sequences, and (iii) complete collection date. I additionally downloaded GISAID reference genome *Wuhan/WIV0J*_*t*_*/2019* (accession id EPI_ISL_402124) for use in the data preprocessing steps. All people involved in sequence data collection are acknowledged in Supp. Mat. D.

Additional removal of sequences was performed on the basis of either one of these two criteria being fulfilled, (i) sequences were not extending from position 250 to 29700 of the reference genome, (ii) the proportion of ambiguously resolved bases was higher than 10%. The remaining 1284 sequences from CH (1673 in DE, 1919 in FR, 1314 in IT) were further classified into 627 alleles (886 in DE, 1166 in FR, 750 in IT) using a custom pipeline available as part of the provided code and detailed in Supp. Mat. A.

### 5.2 MCMC specifications

The timeline of the analysis is fixed for N and S. It extends from the 1st of June 2020 to the 13th of January-2020, cut into successive intervals of 4 weeks each. I assume that *μ* and *g* are known from other studies and are not the focus of the inference. The mutation rate *μ* is fixed to 0.065 mutations per genome per day, corresponding to 8 × 10 ^−4^ mutations per nucleotide per year, and the generation time *g* is fixed to 5 days.

Finally, hyperparameters values for *N* and *S* are chosen so as to not be too informative, using a quick back-of-the-envelope reasoning around Equations (3.5) and (3.6). We imagine what could happen over a time period with few data, as happens in the beginning of the dataset. If the order of magnitude of *N* is approximately 10^4^, and we roughly believe that one out of 5 × 10^3^ individuals is sequenced in reality, because each individual lives for 5 days in the model, it corresponds to sequencing one out of 2.5 × 10^4^ individuals, i.e. *S ∼* 4 × 10 ^−5^. If we now imagine observing this situation over a time period Δ ∼ 10 days, we would observe on expectation 4 births. Using Equation (3.5) helps fix the prior for 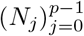 values, which are *𝒢ℐ𝒢* distributed with hyperparameters (*λ, χ, ψ*) = (4,0, 8 × 10^−4^). Then, using Equation (3.6) helps fix the prior for 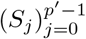 values, which are Gamma distributed with hyperparameters (*α*_*s*_, *β*_*s*_) = (4,10^5^).

I ran the Gibbs sampler for 10^4^ steps, discarded the first 10^3^ steps as burn-in and used the remaining 9 × 10^3^ steps for posterior inference. The code provided along with this article generates the traces and auto-correlation functions of all parameters. These are inspected visually and the ESS values of all parameters are higher than 100.

## 5.3 Results

The output of the Gibbs sampler on each country is a posterior sample of 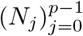 and 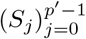 values through time over the fixed timeline. Figure 6 illustrates the input data (to the left) together with the output posteriors of *N* and *S* (to the right). In particular, the trajectories of *N* and *S* tell a similar story in the four countries. The outbreak slowly started in early 2020 and reached its peak around March-April 2020, before quickly decreasing in May 2020. France appear on this Figure to have experienced the longest outbreak. In all four countries, the sampling intensity increased prior to the peak of the epidemic and slowly decreased thereafter.

**Figure 6:**
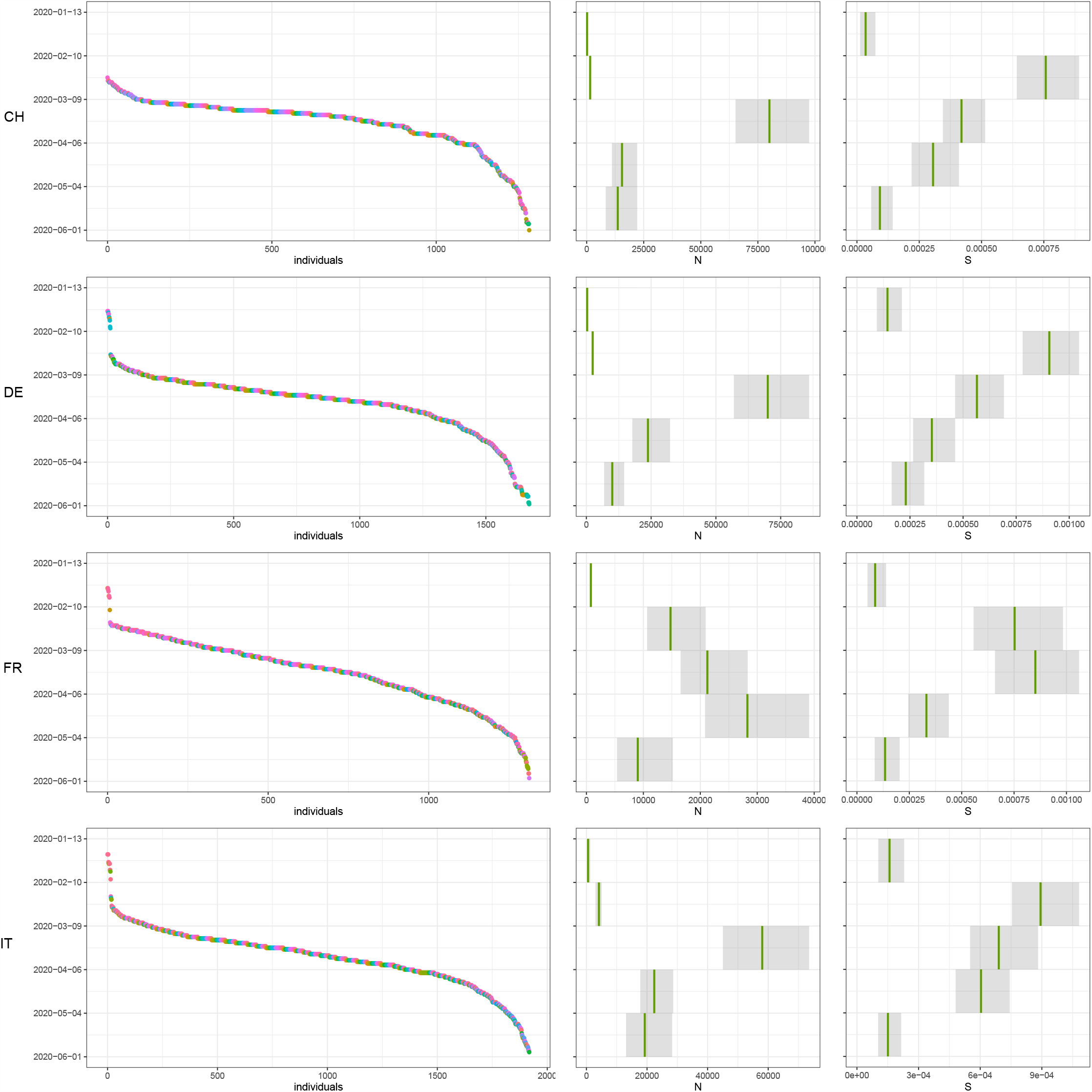
Results of the inference on SARS-CoV-2 data in four European countries during the first wave of the epidemic. CH: Switzerland, DE: Germany, FR: France, IT: Italy. First panel to the left, the raw data consists in an allele partition of sequences sampled through time, where different colors correspond to different alleles. Second and third panels, the posterior sample of *N, S*, with the median shown in green and a 95% posterior interval shown in gray.

**Figure 7:**
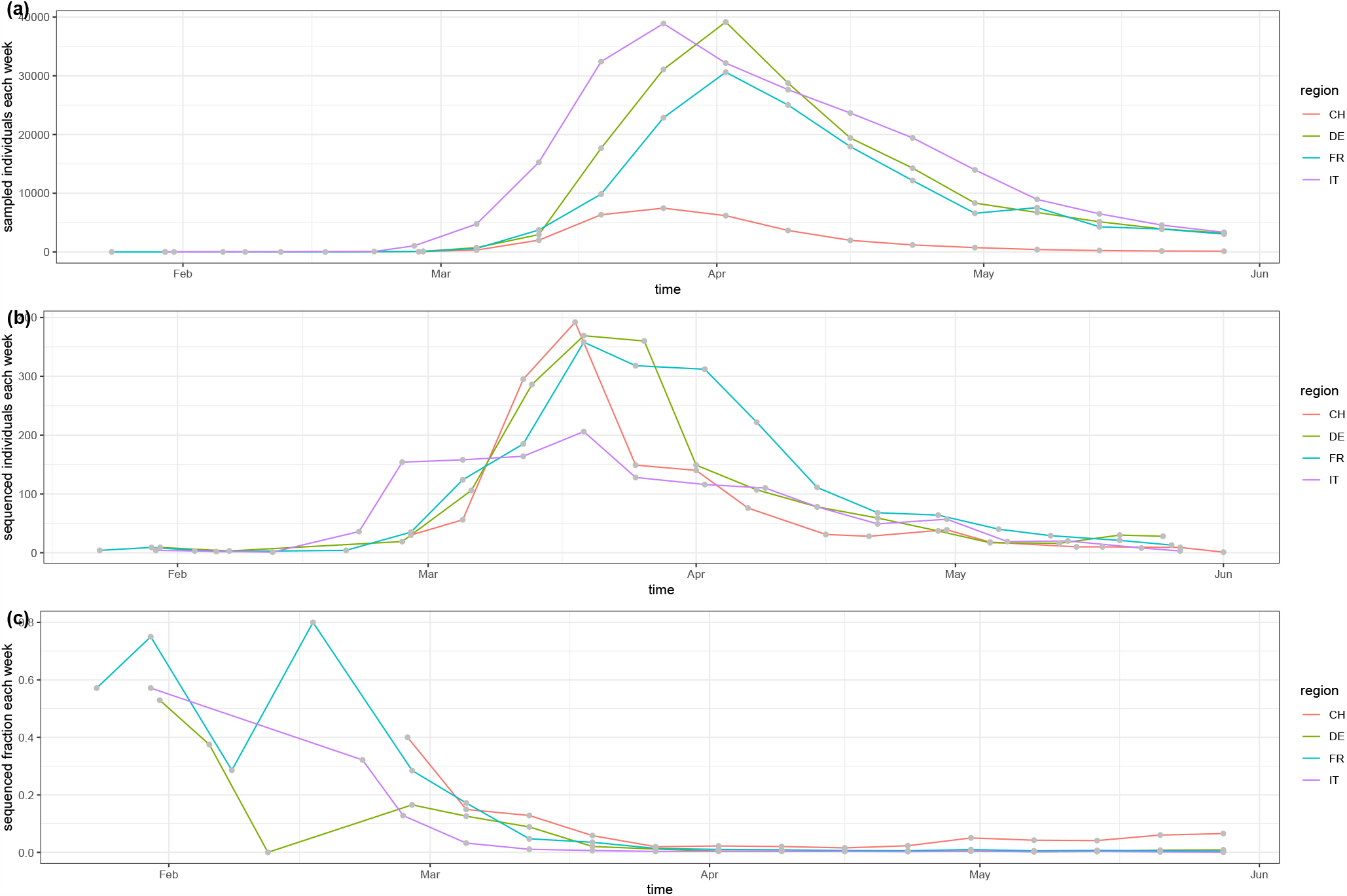
Information on the number of infected, and the number of sequences, sampled through time in CH, DE, FR, and IT.

## 6 Discussion

In this manuscript, I have presented a new Bayesian inference method relying on a Gibbs sampler to sample the posterior distributions of effective population sizes and sampling intensities through time, from the knowledge of an allele partition of sampled-through-time sequences. It relies on a coalescent framework under the infinite alleles model from population genetics, with constant mutation rate, constant generation time, and piecewise-constant effective population sizes and sampling intensities through time. Applying the method to SARS-CoV-2 sequences sampled throughout the first wave of the outbreak in Europe allows us to obtain estimates of effective population sizes, sampling intensities and mutation rate from genetic data only.

### Inferred dynamics on empirical data

In studies relying on a coalescent model, the usage is to focus on the inferred patterns of change through time rather than on the parameter values per se. The rationale explaining this is that values might be affected in complicated ways by various model misspecifications, among which, for example, having particular distributions of offspring numbers at reproduction, having population structure not taken into account, having a different generation time or mutation rate, or having different priors on *N* and *S*. Yet, as long as these model misspecifications do not change through time, on can trust the dynamics of the *effective* population size *N*_*t*_ to reflect the dynamics of the true population size. Although I have followed this convention so far throughout the manuscript, one can notice here that the orders of magnitude of inferred *effective* values seem nevertheless to roughly match what is known from the case count record only (shown in Supp. Mat. B).

In terms of dynamics as well, the patterns of effective population sizes appear compatible with what is known about the first wave of the outbreak in these countries. The number of infected individuals starts increasing exponentially early in 2020, reached a peak in March-April, before decreasing in response to policy measures taken -different versions of a lockdown have been implemented in the four countries around this time. Interestingly, the sampling intensity, on the other hand, starts increasing earlier during the epidemic, when the research effort started growing around SARS-CoV-2 genetics but the case counts where not so high yet. It then decreased again following the peak of the outbreak.

### Opening future modeling opportunities with allele data

Overall, the results on empirical data illustrates the use and applicability of the method on large real-world datasets. Yet, the current model formulation still lacks a few realistic ingredients before it can be used to learn new aspects of an epidemic.

First and foremost, it does not take into account population structure, a feature that is likely to be present in the empirical dataset and violates the model assumptions. Further work is needed to properly incorporate different demes characterized by different population sizes and sampling intensities, exchanging sequences through migrations among demes. This will likely be the subject of future work aiming to infer population structure and population dynamics using the allele partition. Integrating over the hidden coalescent history will be complicated a bit by migrations between demes, but could be envisioned within a closely related inference framework.

Second, it could be interesting to use smoothing priors for *N* and *S* to ensure that these two functions of time do not show huge steps from a time period to the next. This will as well be the focus of future developments of the method, that could build on related work in phylodynamics (Karcher et al. 2020; Parag et al. 2021). It would seem especially interesting for *N* to root smoothing priors on mechanistic assumptions of population dynamics, such as considered e.g. in other epidemiological models (Cori et al. 2013).

Last, the boundaries of the intervals on which *N* and *S* remain constant could be unknown, with or without prior distribution. This would allow the timeline to be informed by the data, and could lead as well, in case it is stochastic and averaged over the posterior, to smoother N and *S* trajectories. This could build on related work in a phylodynamic context under a birth-death model (Stadler 2011).

Further theoretical work is also required before this kind of model can be applied to larger -already available -datasets. Indeed, while the initial hope of bringing back into fashion the infinite allele model was to be able to process very large genetic datasets, it is in the end not realistic to use the current implementation of the method with more than ∼ 10^4^ sequences. At least four directions can be envisioned to improve on this objective in the future: (i) trying to optimize the sampling step of the coalescent history, which is the current bottleneck of the computation, so that it scales closer to *O(B)* than *O*(*B*^2^), possibly using well designed approximations; (ii) turning to a maximum likelihood framework, possibly relying on an expectation-maximization algorithm to either completely bypass the need for posterior sampling through MCMC, or at least use it to drastically speed-up the burn-in of the MCMC; (iii) turning to a variational Bayes method properly designed to sample the posterior in our model; and (iv) optimizing the implementation of Algorithms 1 and 2, where the most straightforward idea would be to parallelize MCMC chains for example.

### When is the use of a simplified mutation process pertinent ?

A natural question arises when thinking about the difference between this method and currently used coalescent-based method in phylodynamics, namely *How do these compare in terms of statistical power ?* Or to rephrase it in more technical terms: *what signal do we loose by forgetting about the coalescent history above the first mutation ?*

The answer will likely depend on the mutation rate and on the temporal scale that one is interested in studying. In the limit of very high mutation rate compared to the temporal scale under study, only singletons are observed, bearing no useful signal, while in the limit of very low mutation rate, only one allele is observed, again bearing no useful signal to infer *N* and *S*.

In between, there is a setting with an intermediate mutation rate, such that alleles extend for some time across the focal time-frame, providing signal to reconstruct N and S. However, even in this optimal setting for the method, data is being discarded so that all signal on the internal branches linking different alleles is lost as compared to coalescent-based methods integrating over the full unknown tree. When does the trade-off between computation time and precision turn in favor of using a simpler infinite alleles model ? Quantifying this more precisely on simulations would be a valuable contribution.

Moreover, when one is not interested in estimating p,, the allele partition could be chosen so as to tend to an optimal setting as described above. In principle, the allele partition of the set of sequences could be obtained by applying any another equivalence relation, e.g. *being similar only on a given subsequence*, or *having a similar amino-acid sequence*. These could be used to decrease the number of alleles in the dataset.

In between the two above-mentioned extremes of (i) using an infinite alleles model or (ii) using a finite site model with a substitution model, lies also the opportunity to revive another assumption from population genetics, namely the *infinite sites model*. Under this model, each mutation hits a new site along the sequence, and thus a more precise phylogenetic history between sequences can be reconstructed. An *Importance Sampler* algorithm has been proposed by Stephens and Donnelly (2000) for simulating the past history under a coalescent with an infinite sites model. A more thorough comparison of this inference framework against another inference method relying on the infinite sites model could as well be a relevant contribution to the field.

Finally, this manuscript also opens the way to develop better approximations aiming at taking into accounts more sequences in scenarios with high numbers of duplicates (Boskova and Stadler 2020). This line of research could benefit from the joint use of different mutation models clearly distinguished in distinct parts of the evolutionary tree of sequences, while still relying on a unique underlying population dynamics model, such as e.g. a coalescent model with discrete population size shifts.

### Conclusion

Bringing back into fashion old population genetics simplifications of the mutation process and incorporating them into modern statistical frameworks could play a key role in better surveying and understanding population demographics and structure from molecular data. I hope that this work will participate in a current trend towards adapting computer-intensive phylodynamics methods for use with datasets characterized by low genetic diversity such as the current SARS-CoV-2 outbreak.

## Acknowledgments

This work was supported by an ETH Zürich Postdoctoral Fellowship. I thank the cEvo group at the D-BSSE department for interesting discussions on phylodynamics methods.

## A Details on the numerical method

### Pipeline for creating the allele partition

Creating the allele partition is a central step in the method, for this will be the raw data taken as input by the Gibbs sampler. To build it, I relied on the following pipeline:

1. cut the master reference sequence from position 250 to 29700, and put this one as the first element of a list of reference sequences.
2. initialize an empty list of alleles.
3. Then iterate, for each other sequence, the following steps,
  a. look for a pattern close to the 30 first nucleotides of the master reference into the first 500 nucleotides of each sequence, to get the beginning of the window, and discard the very few sequences where the beginning could not be found.
  b. keep as the focal sequence the 29450 nucleotides following the beginning of the window.
  c. record the collection of snps differing in the focal sequence as compared to the reference sequence.
    - if it is smaller than 100 snps long, we consider it to be the *compressed representation of the sequence* against this reference.
    - it not, compare to the next reference sequence in the list.
  d. if no reference sequence is less than 100 snps different than the sequence, it is likely to have a feature of its own (typically, a gap). Add this sequence to the list of reference sequences.
  e. Look for the same compressed representation of the sequence in the list of alleles and add the sampling date of this sequence to the list of sampling dates of the allele.

This pipeline is implemented in the script `raw_to_datasets .ml`, available as part of the code associated to this article.

### MCMC output analysis

MCMC samples output by the *Ocaml* code are then analyzed using scripts written in the *R* programming language. Post-processing steps rely in particular on the following packages to work,

- the very versatile *ggplot2* and *cowplot* to produce figures,
- *LaplacesDemon* to compute ESS values using traces of scalar values,
- *forecast* to compute ACF or PACF using traces of scalar values,

The R post-processing scripts are also available as part of the code released along this article.

### Generation of random variables

Random variables distributed according to a 𝒢 ℐ𝒢 distribution are sampled using a personal implementation of Hermann and Lcydold (2014)’s algorithm in the programming language *Ocaml*. It is naive translation of the very handy R package *GIGrvg* by the same authors.

## B Case count data in the four countries

Case count numbers in the four countries of interest have been downloaded from the WHO website on the 16th of June 2021. I plot here the number of samples, number of sequences, and sampling fraction through time over the time frame of the first wave of the outbreak, for comparison with the order of magnitudes of *N* and *S* presented in the main text.

Note here that the *sequencing fraction* plotted below corresponds to the number of sequences divided by the sum of the number of sequences and number of samples. Indeed, the number of sequences in the early time period is sometimes higher than the number of samples, and I thus considered that the sequences were not included in the case count.

## C Gibbs sampler code documentation

This Section aims at making the code released along this article more easily comprehensible, by shortly presenting in plain text the strategy to store different data, together with key functions to manipulate the data.

### Format of key quantities

By default, all lists are ordered in time ascending order (that is, first element on top of the list at present time 0).

`lists` **or** `listN` list of ordered (t_*j*_, S_*j*_), where *S*_*j*_ is valid on [*t*_*j*_, *t*_*j*+1_).

**an** `allele` **or a** `lineage` an ordered list of either *(b*_*i*_) belonging to the same allele/lineage, if the individuals are not numbered, or an ordered list of (*n*_*i*_) if the individuals have been numbered.

`alleles` **or** `lineages` a list of (ordered lists of (*b*_*i*_)) or a list of (ordered lists of (*n*_*i*_)) depending again on whether individuals are numbered or not.

`array_individuals` an array of information about individuals, with element number *i* referring to individual number *i*. Each element is a tuple *(b*_*i*_, *h*_*i*_, *o*_*i*_, *a*_*i*_) where b_*i*_ is the birth time, *d*_*i*_ is the death time, *o*_*i*_ is the number of the individual into which this individual coalesces (or *o*_*i*_ = *i* if there is a mutation), and *a*_*i*_ is the ID number of the allele this individual belongs to.

`array_alleles` an array of information about alleles. Each element number *a* is the list of ID numbers of all individuals belonging to allele *a*.

`samp_history` an ordered list of (*b*_*i*_, *i*) where *i* is the ID number of the newly born individual at time b_i_.

`coal_history` an ordered list of (*h*_*i*_, *i, o*_*i*_) where *i* is the ID number of the individual dying at time h_i_ by coalescence into lineage *o*_*i*_ (or by a mutation if *o*_*i*_ *= i)*.

`all_events` a list of all events ordered in time ascending order, with lots of other interesting quantities attached to the interval between this event and the following, such as : the number of lineages alive, the current values of *N*, the current value of *S*, the total rate at which an event happens on this interval for a given individual, etc…

`intervals` the ordered in DESCENDING order list of 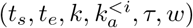 where *t*_*s*_ is the time of start, *t*_*e*_ is the end time, *k* is the total number of lineages alive on the interval, 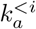 is the number of individuals from allele *a, to* the left of individual *i*, being alive on the interval, *τ* is the total rate at which death happens and *w* is the weight of the interval, i.e. the probability that the death event of the focal individual falls in this interval, conditioned on *𝒜*. Note that the reason it is in descending order is that it is built by reading the list of `all_events` in ascending order, and the next steps consisting in drawing the interval does not require any specific order.

### Roles of some key functions

Here is now an overview of some of the key operations we need to perform for Gibbs sampling steps. **simulation if needed** to get `samp_history`

first using sim_sampling_events.

And then to get alleles using `sim_coal.`

**pre-processing** we build the `array.individuals` and `array.alleles` from the alleles and samp-history.

**Gibbs sampling** We need to consider in turn the following operations,

- Updating the death time of individual number *i* using `update_past_coal:` requires to first build the weighted intervals with `get_intervals`,before drawing an interval with `find_back_interval`, simulating the death time within the interval and finally erasing the old time and inserting the new at a correct place in `all.events` using `replace.coal`.
- Updating the *(N*_*j*_) values using `update_listN`, which explores the list of `all.events`, and modifies in place the *N*_*j*_ values.
- Updating the *(S*_*j*_) values using `update_listS`, working similarly as above.
- Updating *μ*, using `update_imi.`

## D Acknowledgements for data collection

I am grateful to the following list of authors, who have contributed to the collection of SARS-CoV-2 sequences that I downloaded on GISAID. In the four tables below, you will find the authors that took part in the collection effort in, respectively, Switzerland, Germany, France and Italy over the time period I have been interested in.

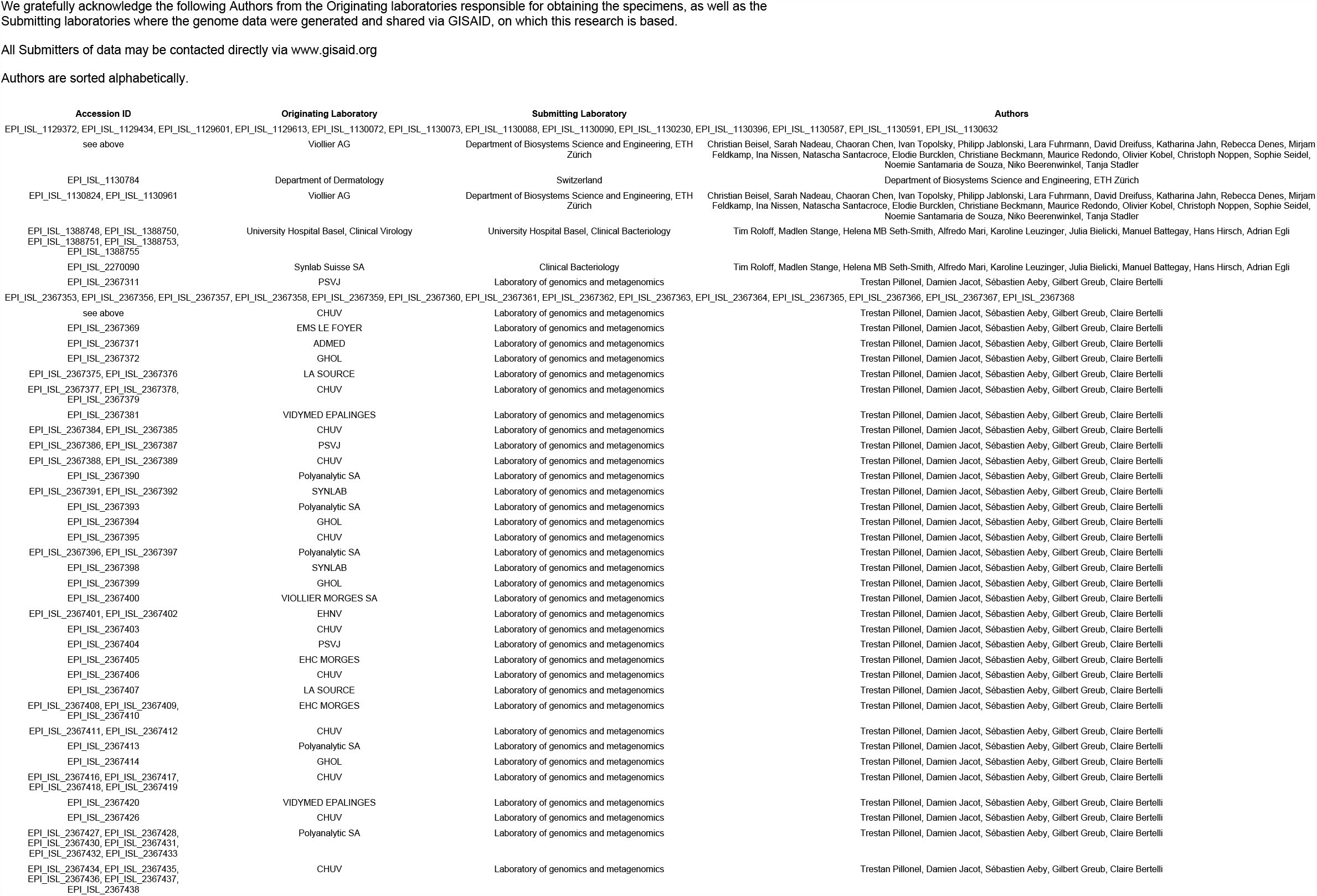
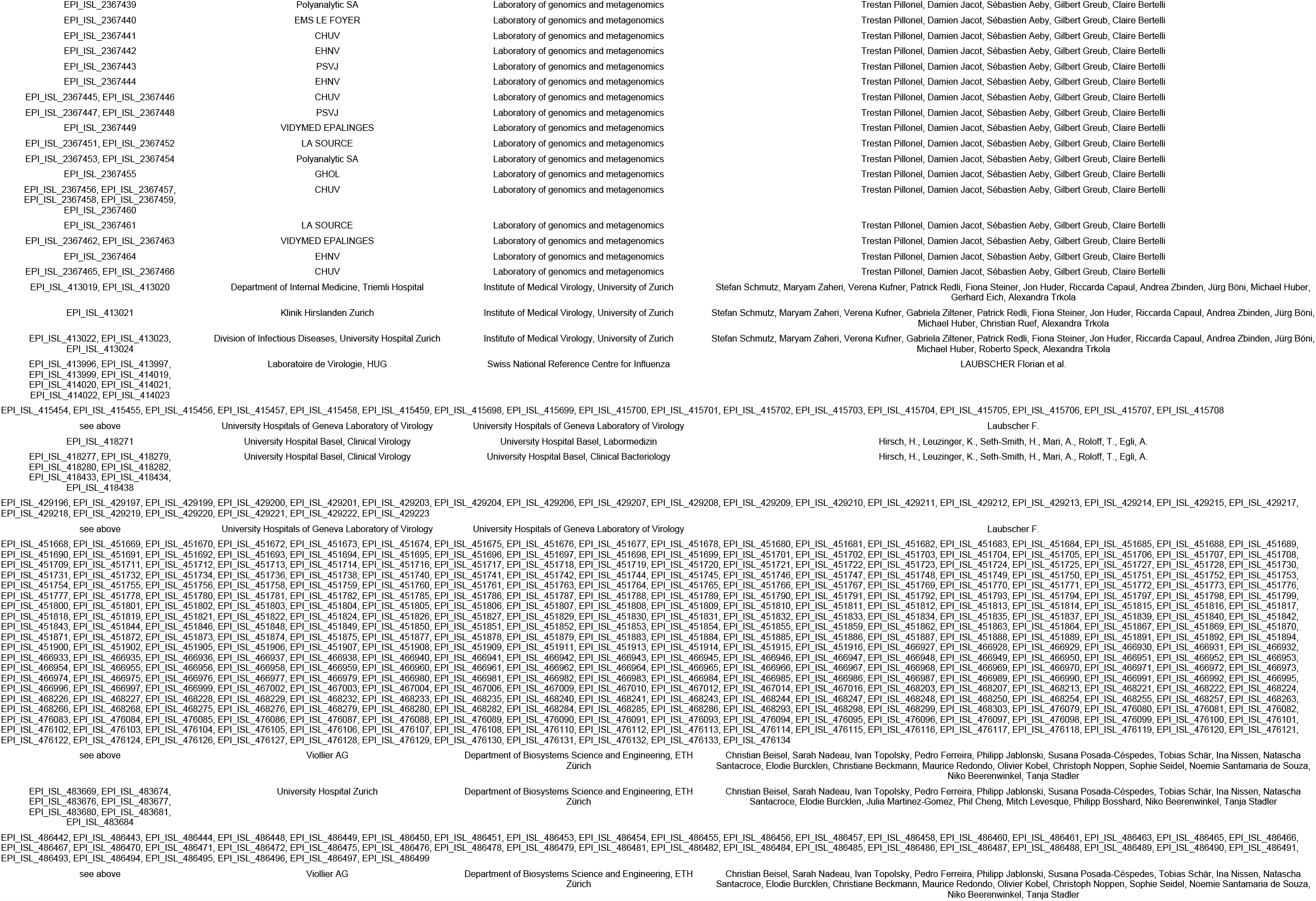
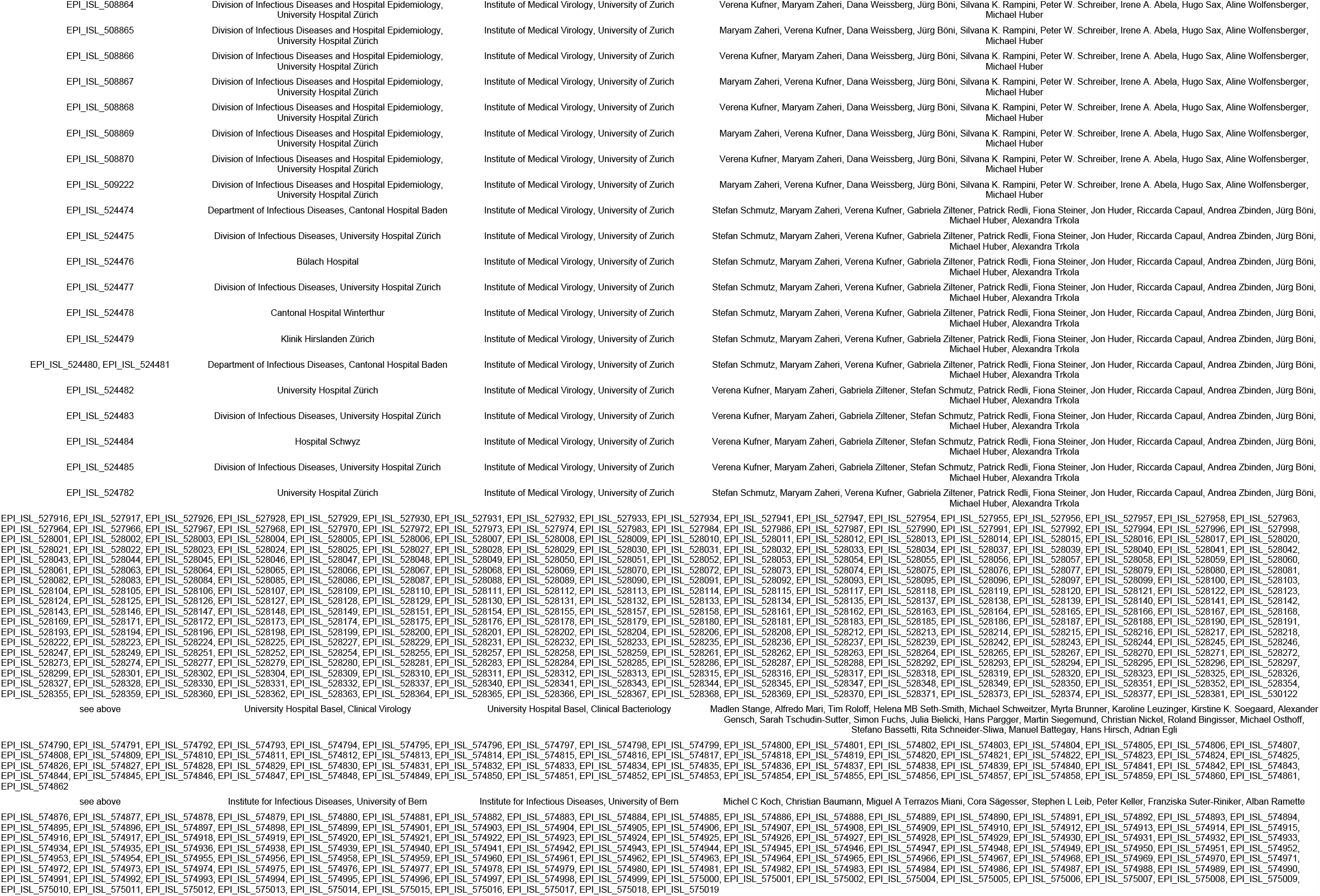
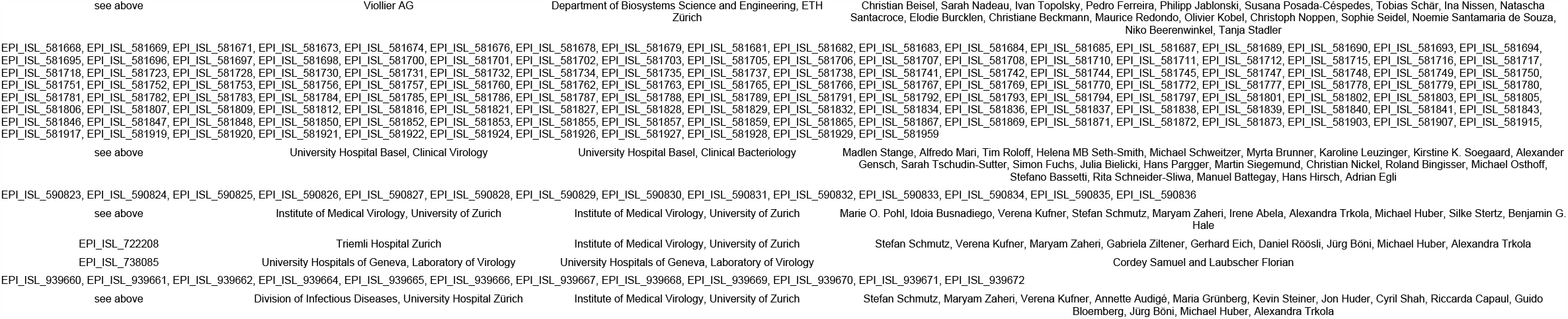
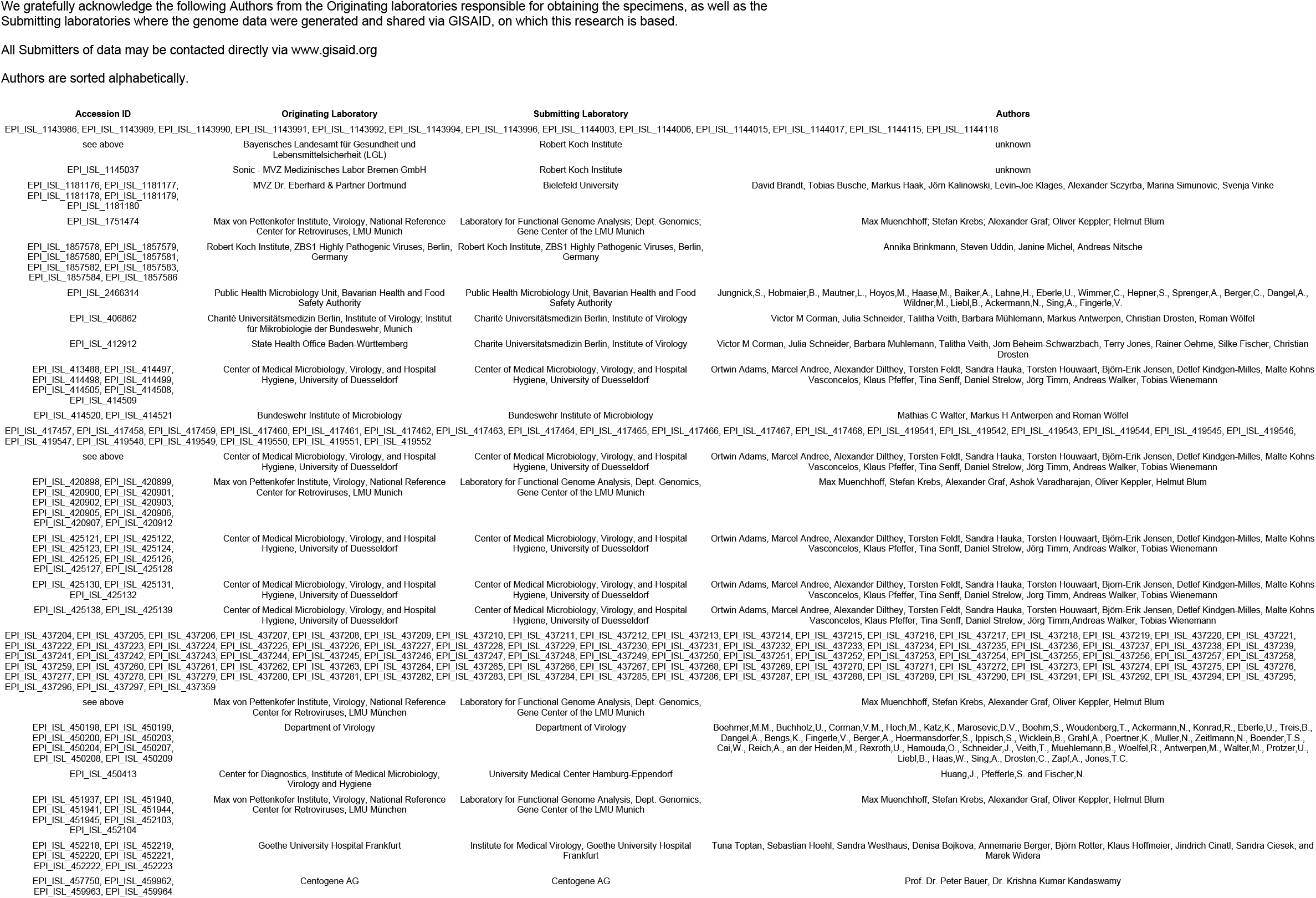
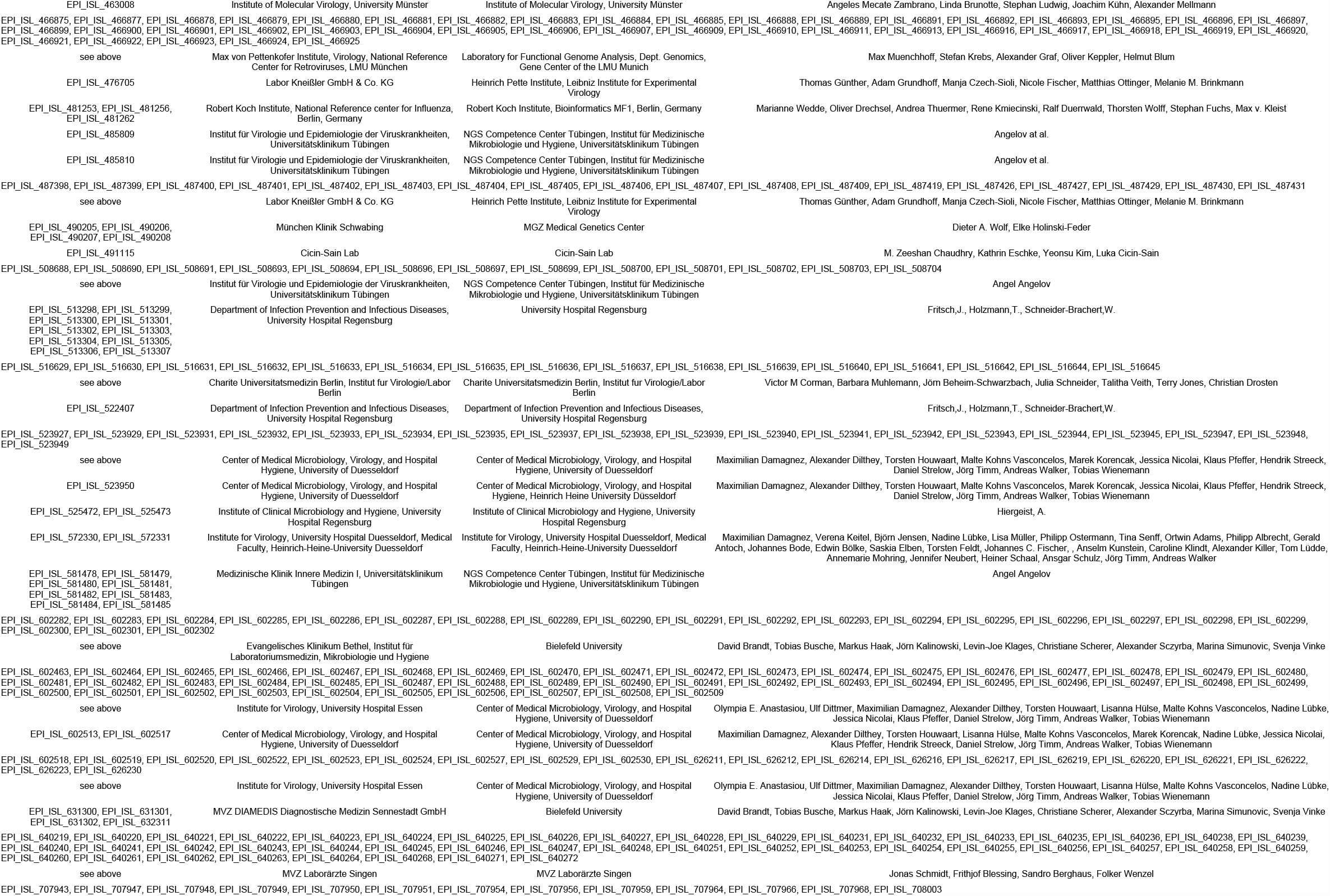
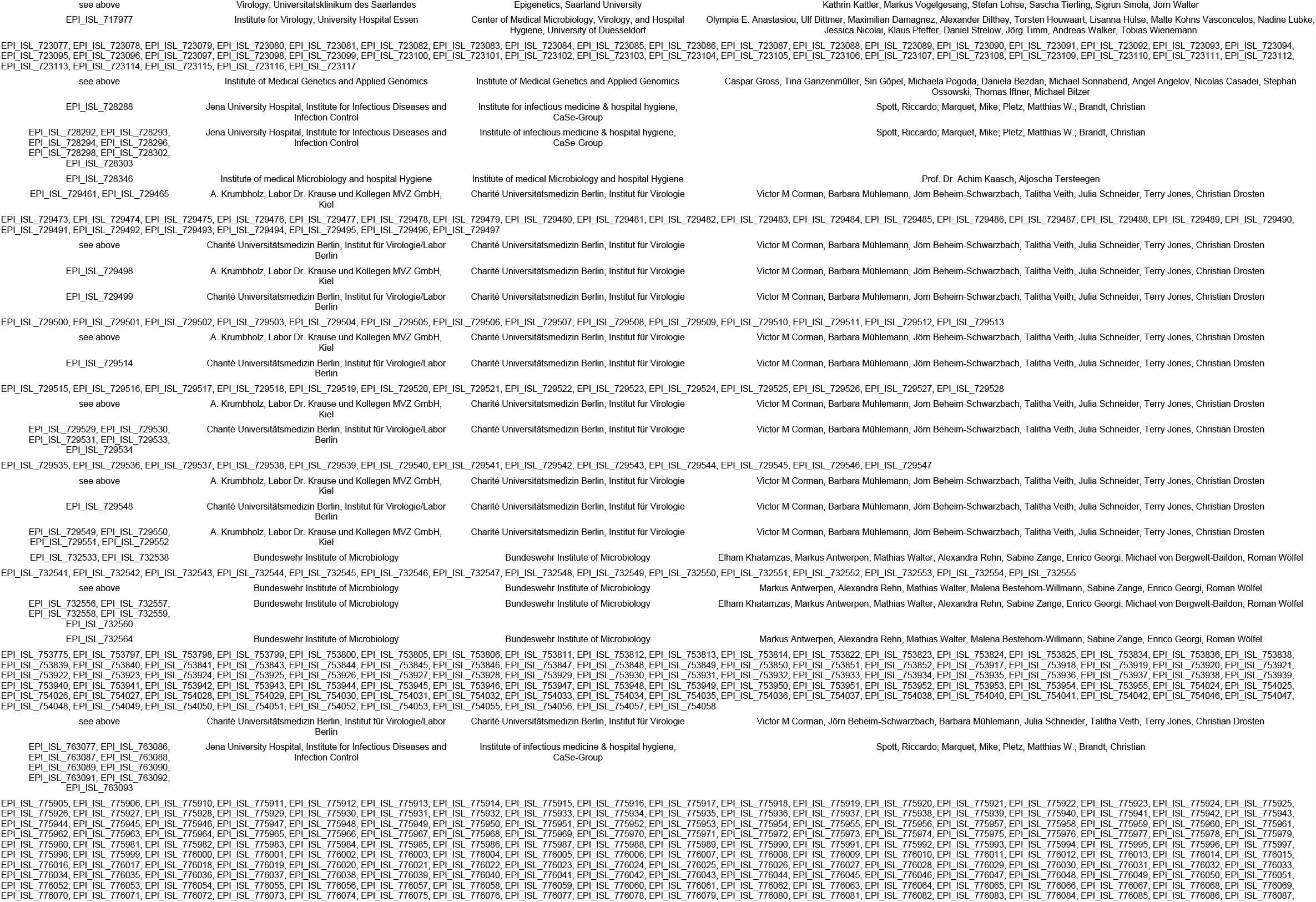
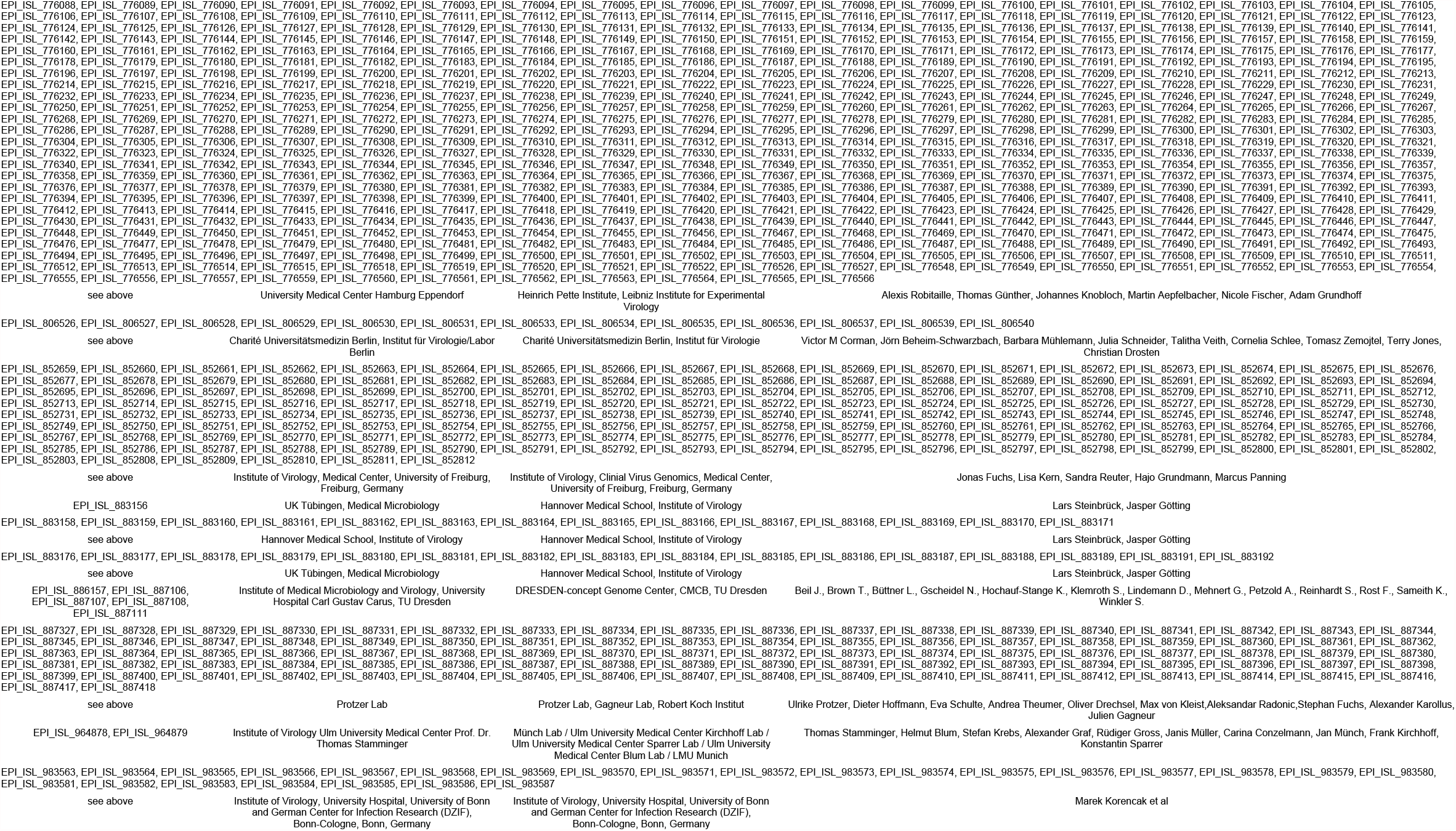
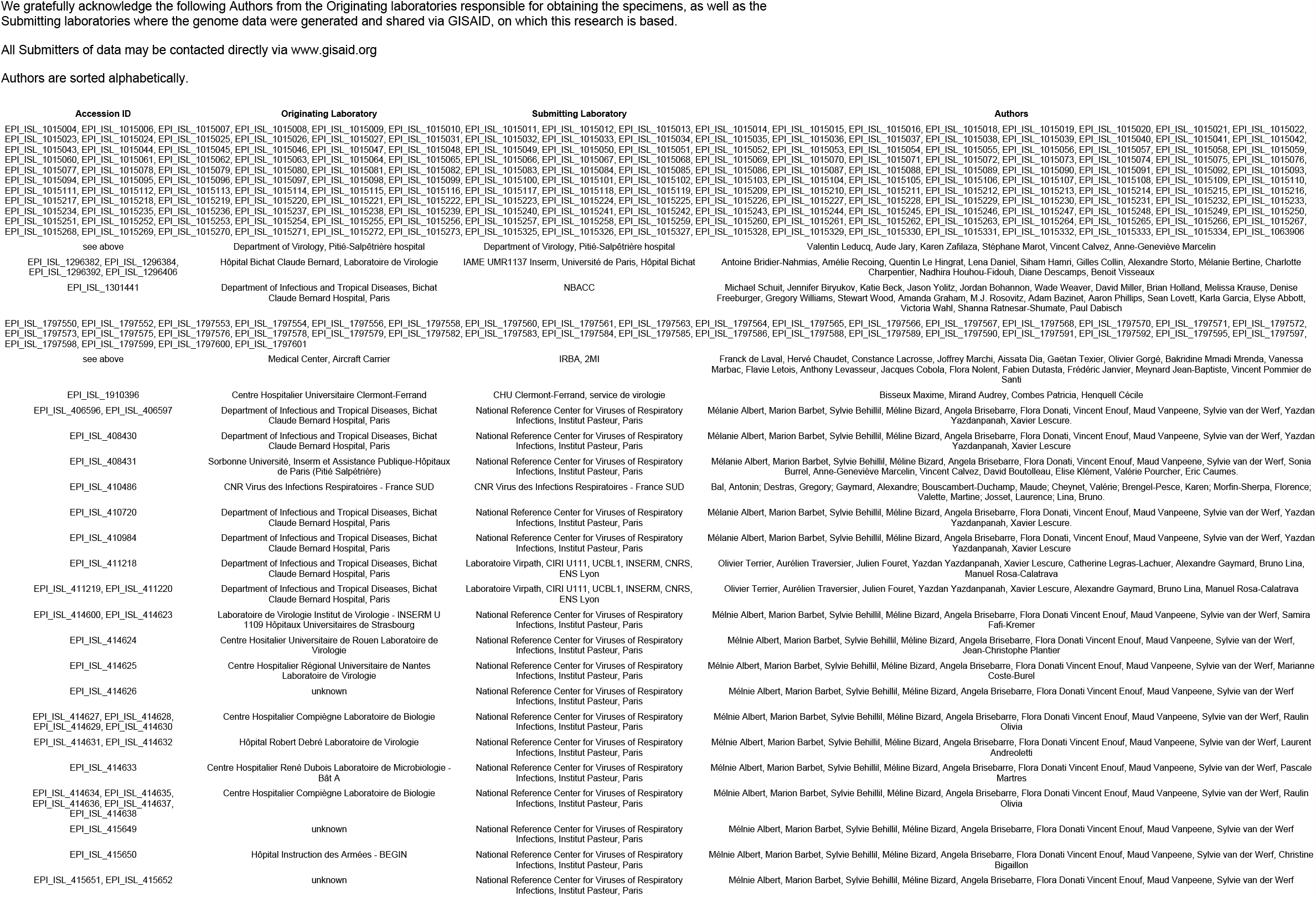
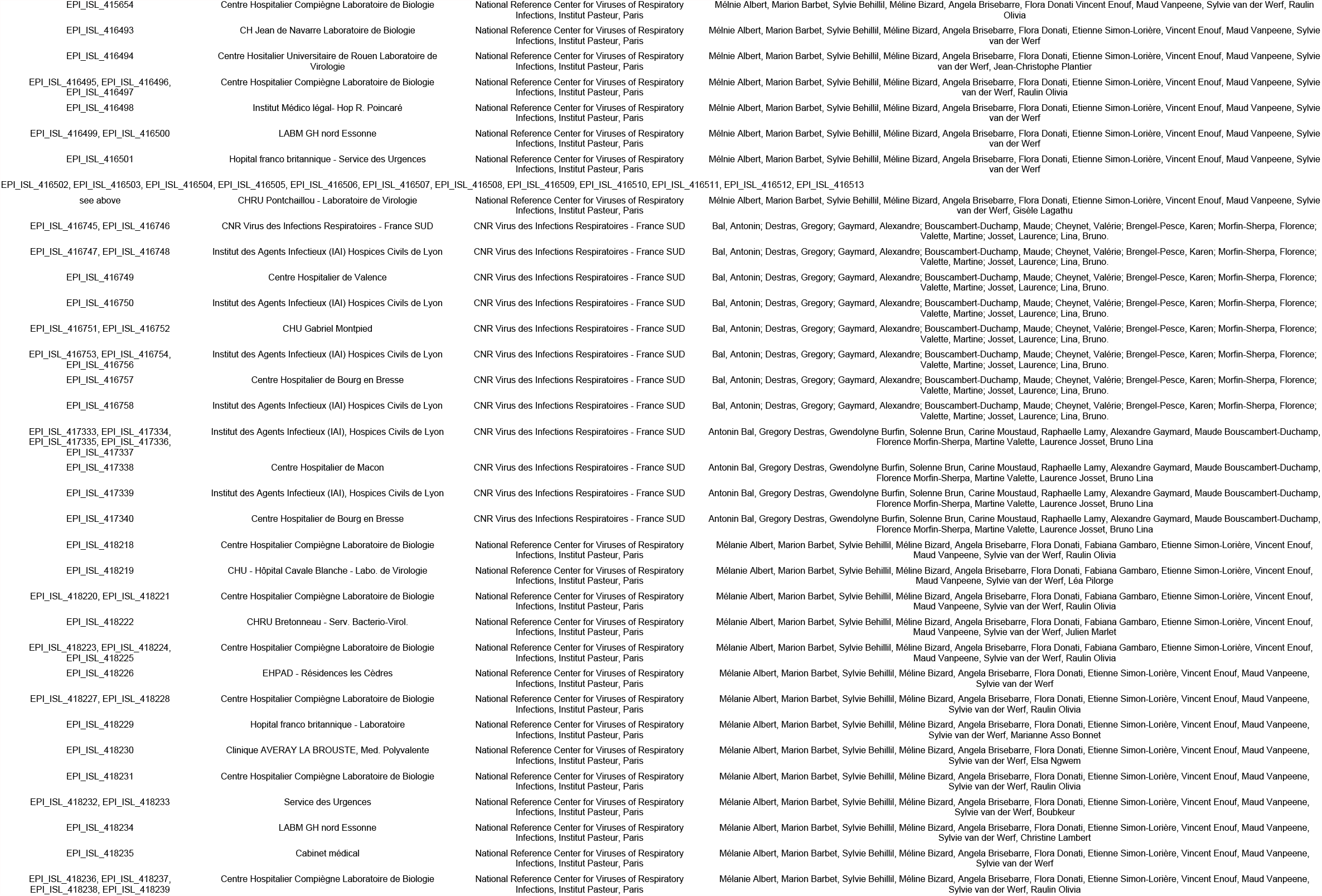
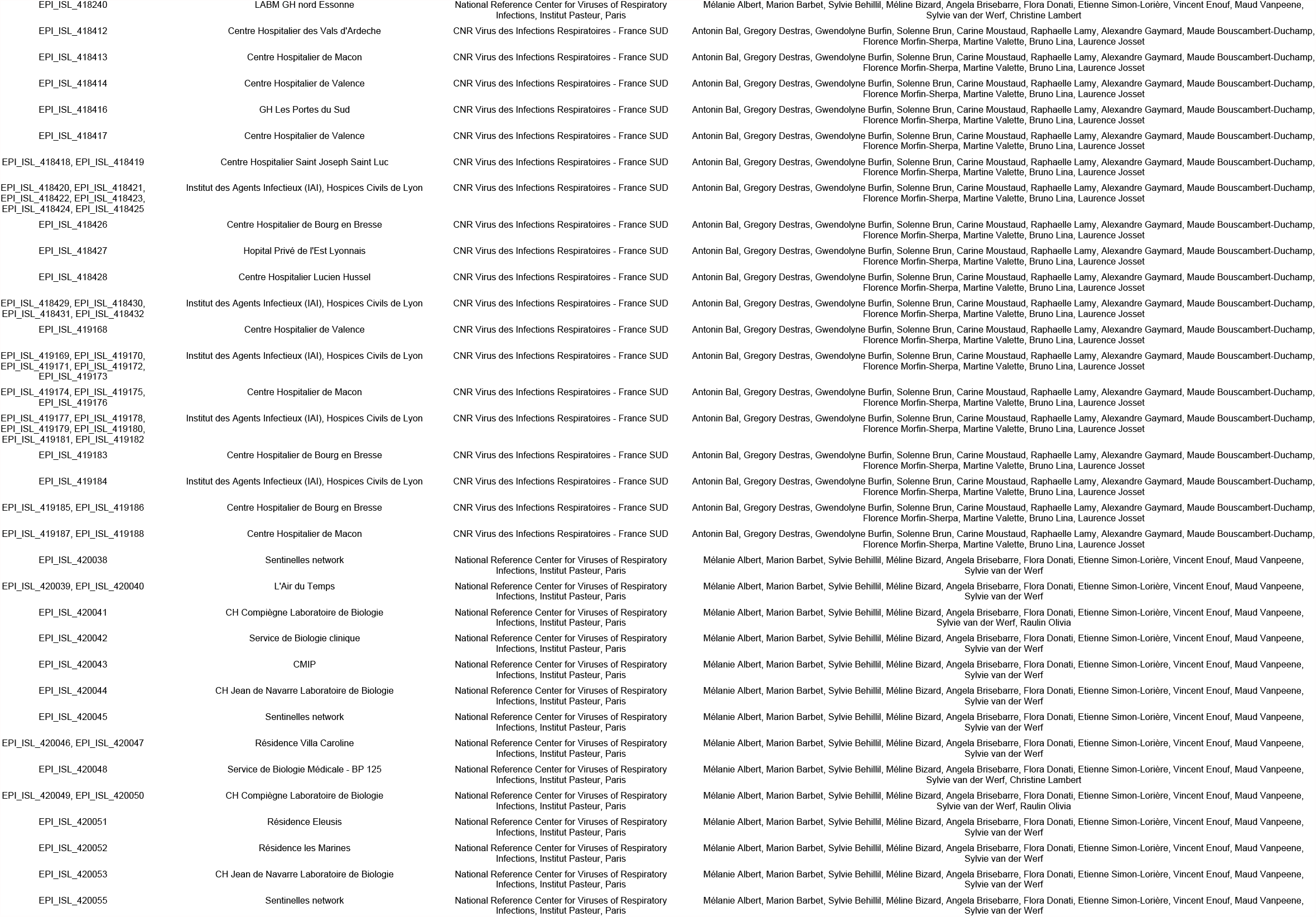
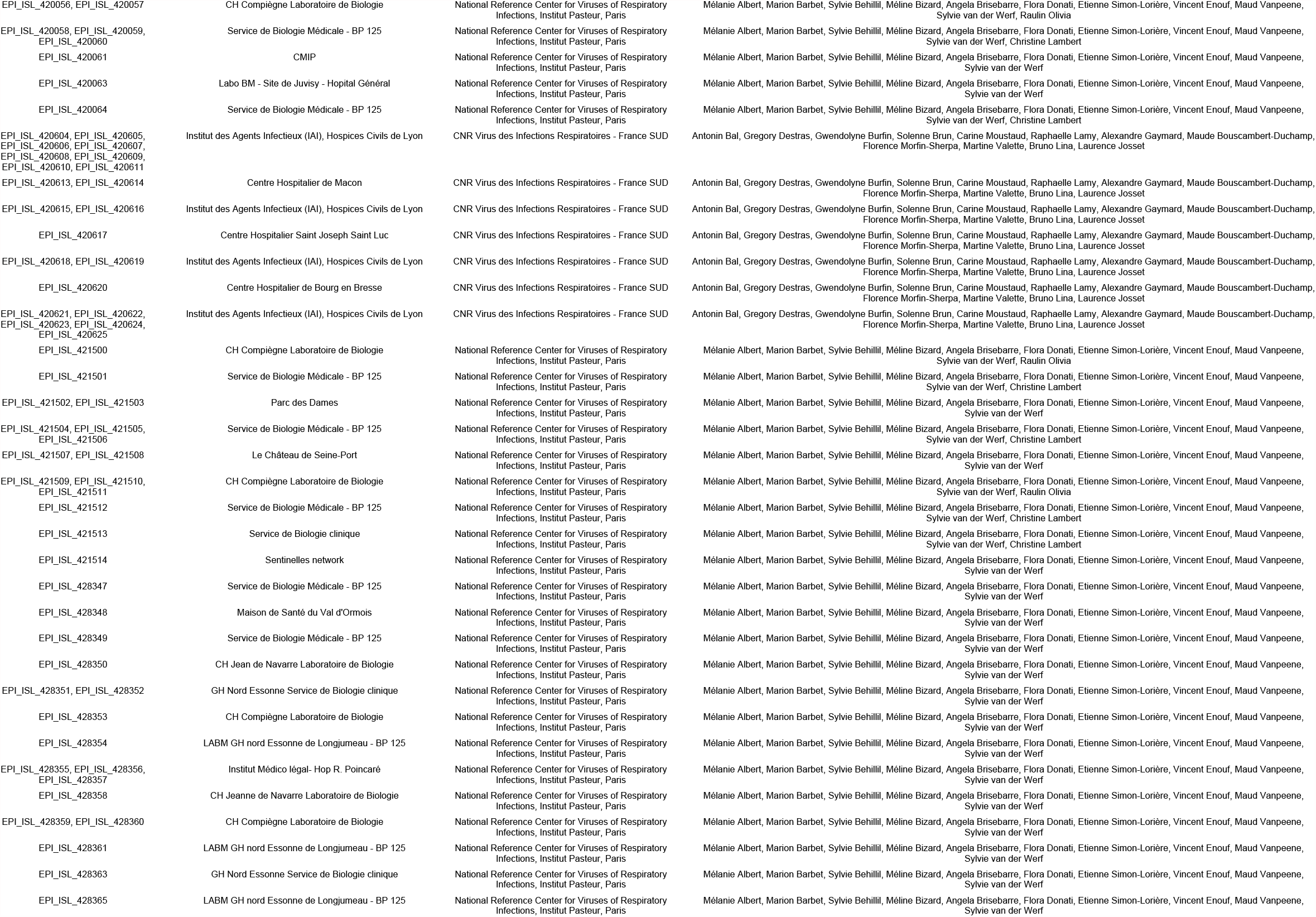
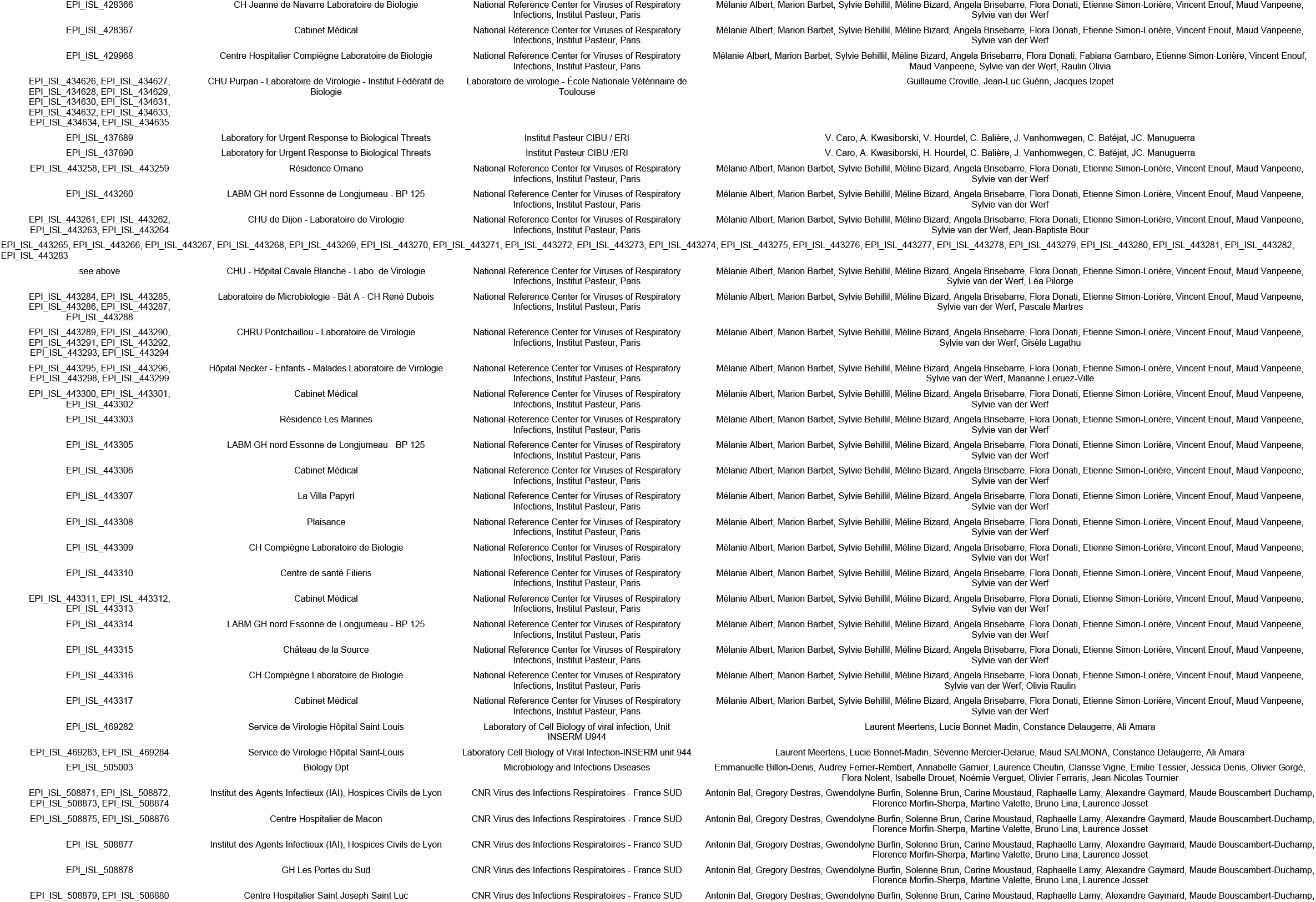
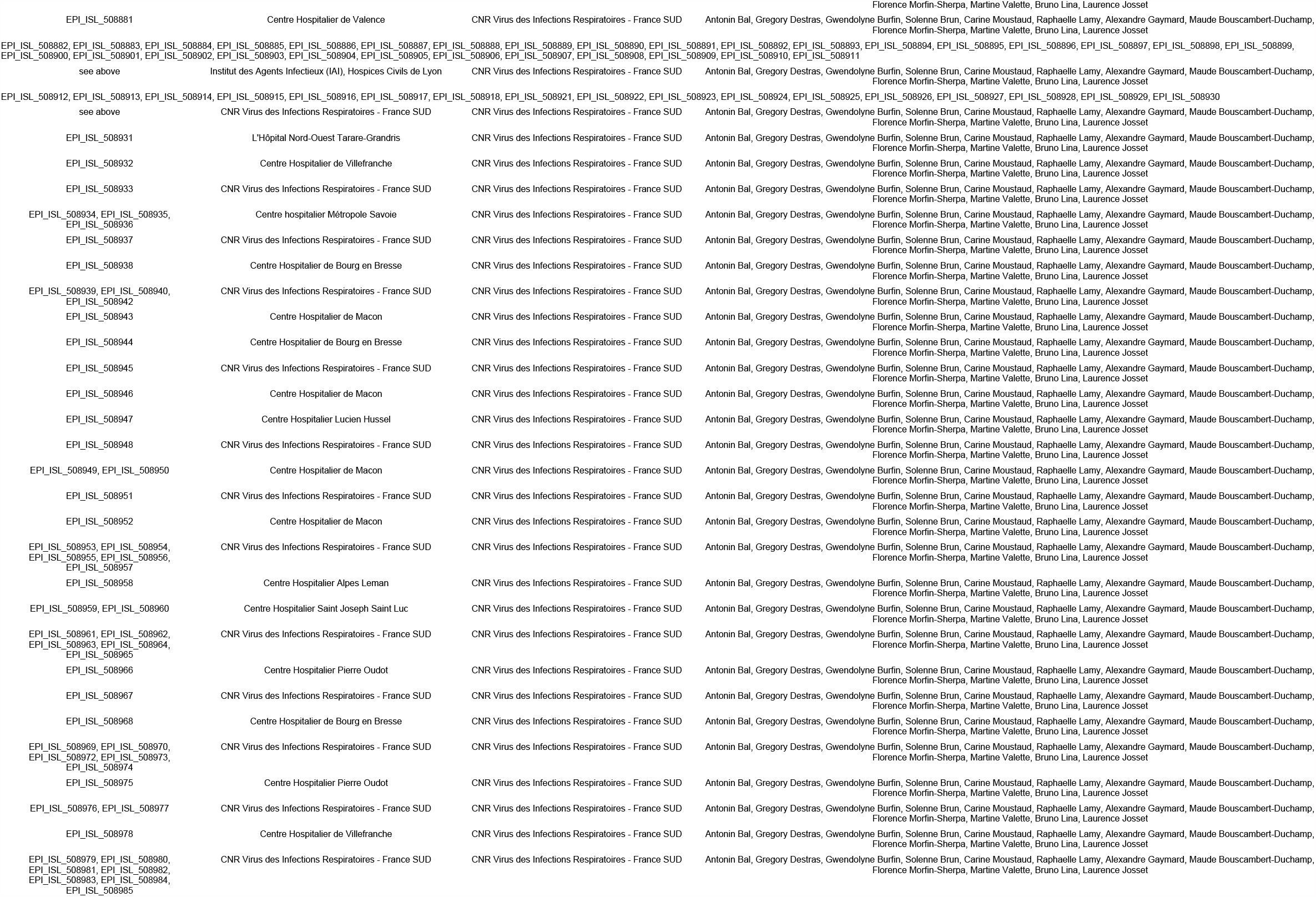
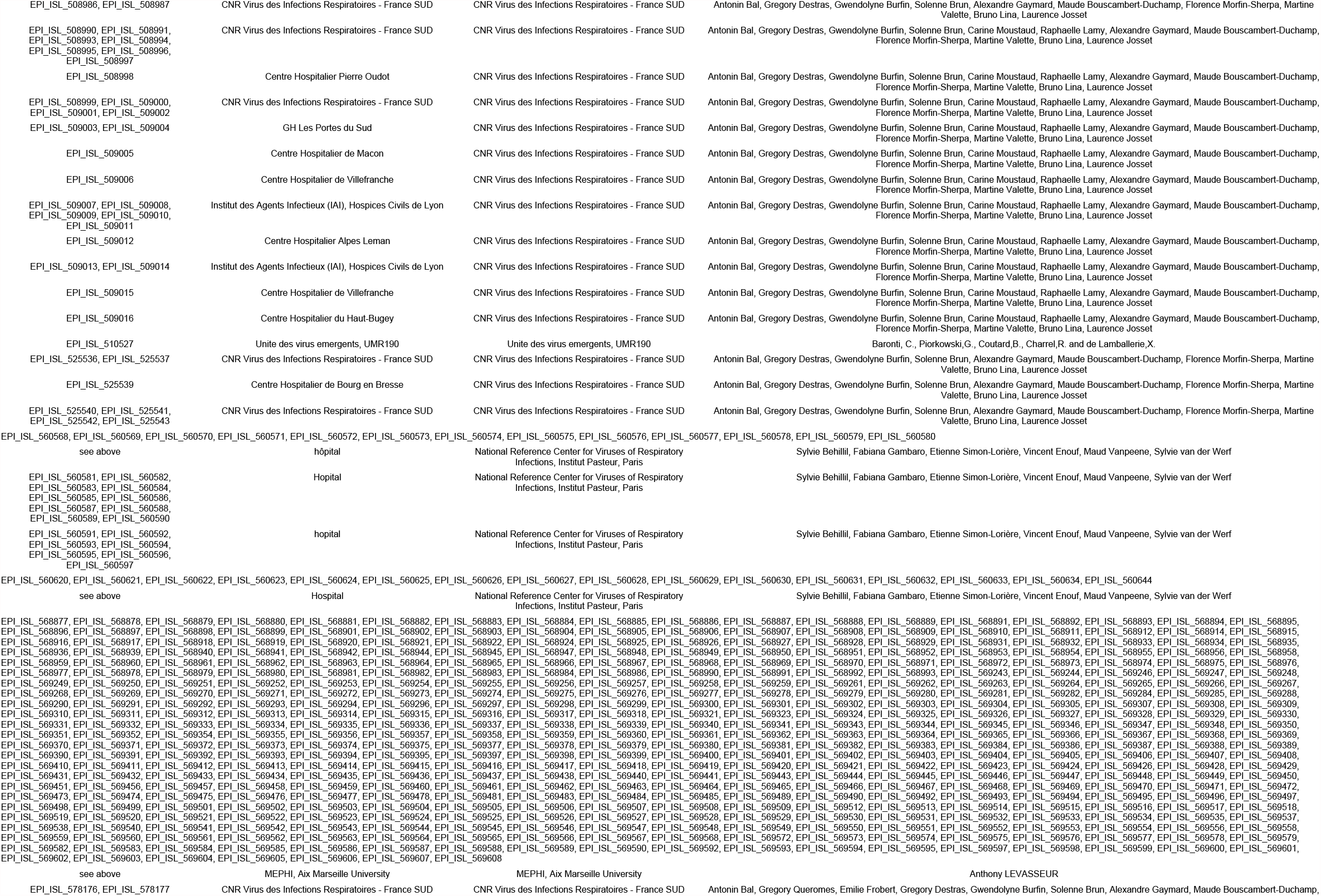
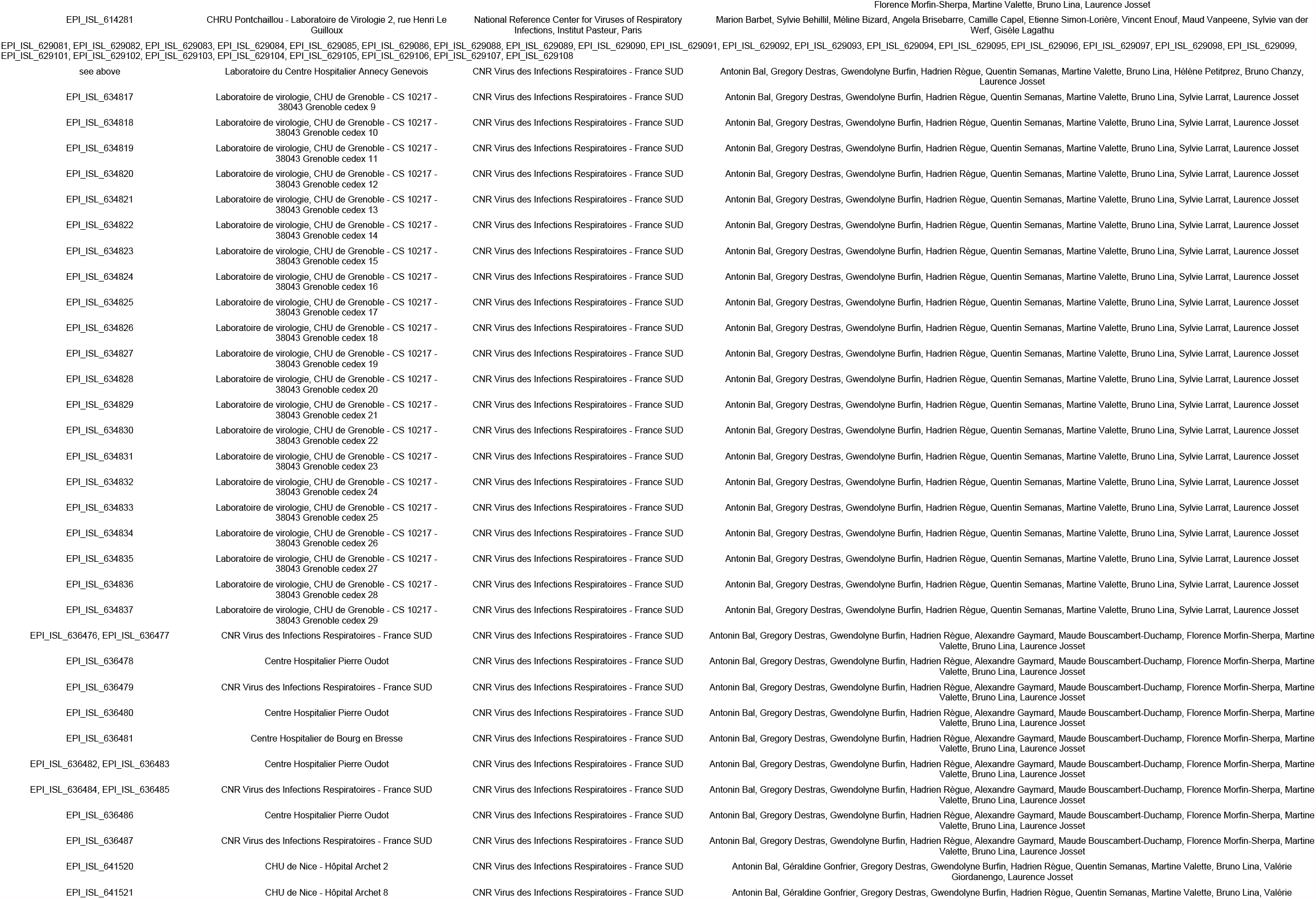
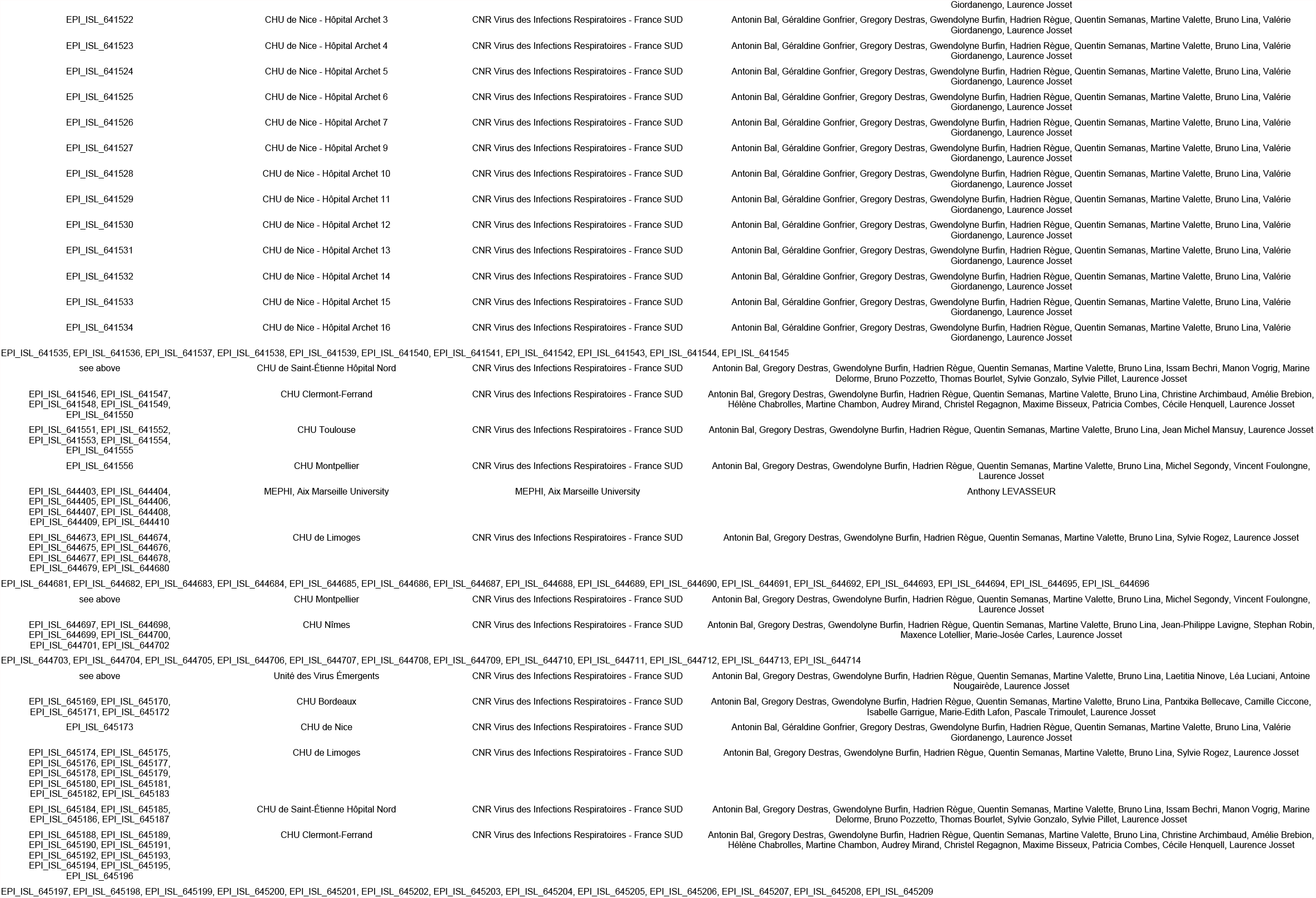
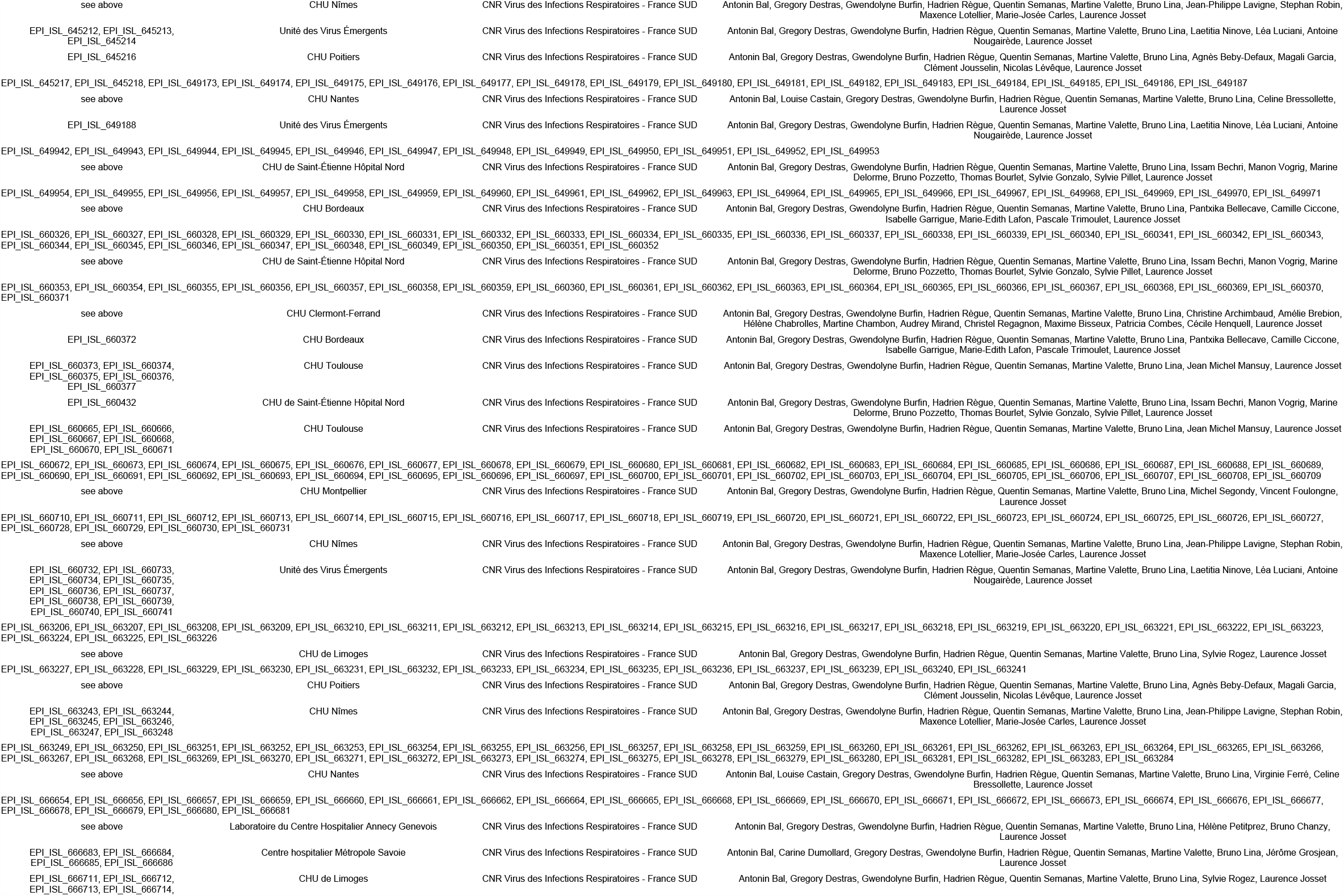
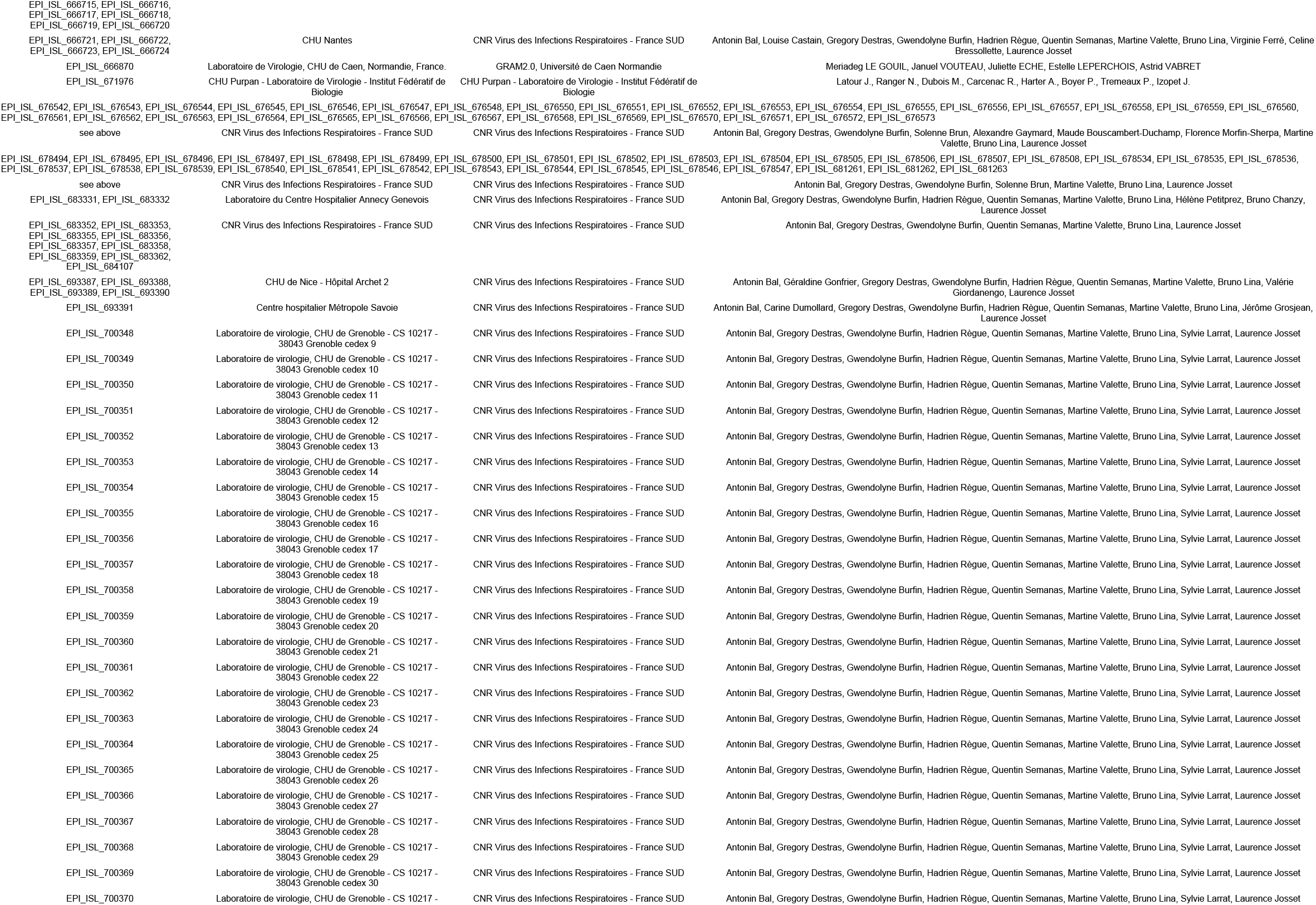
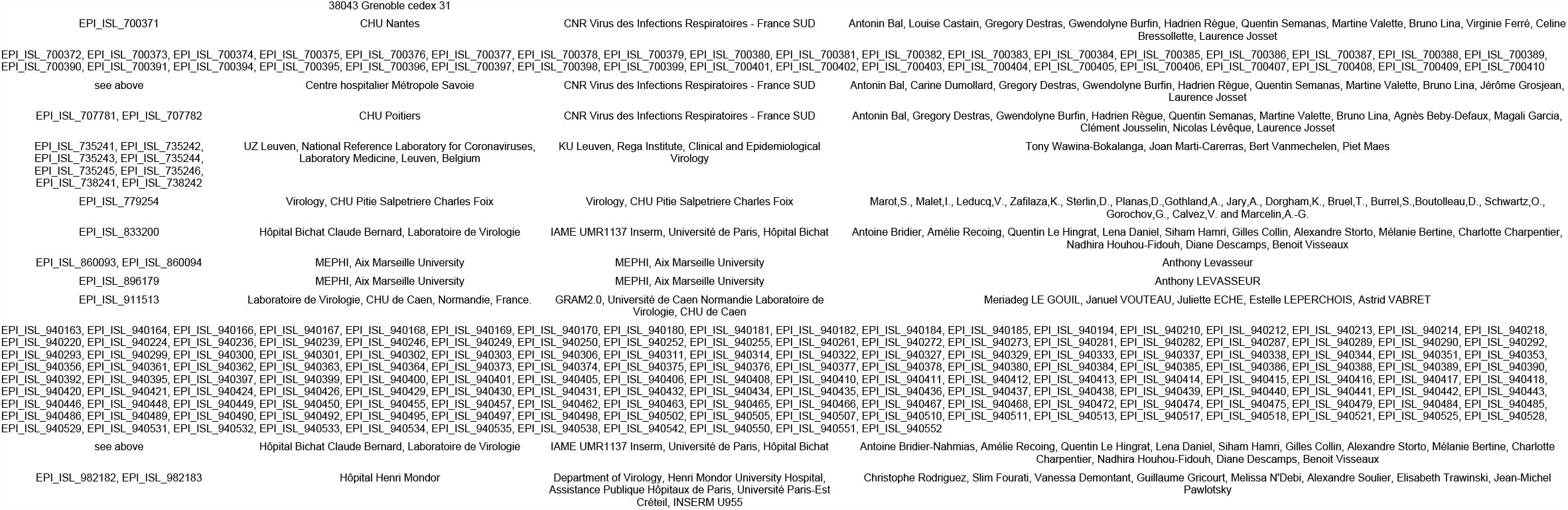
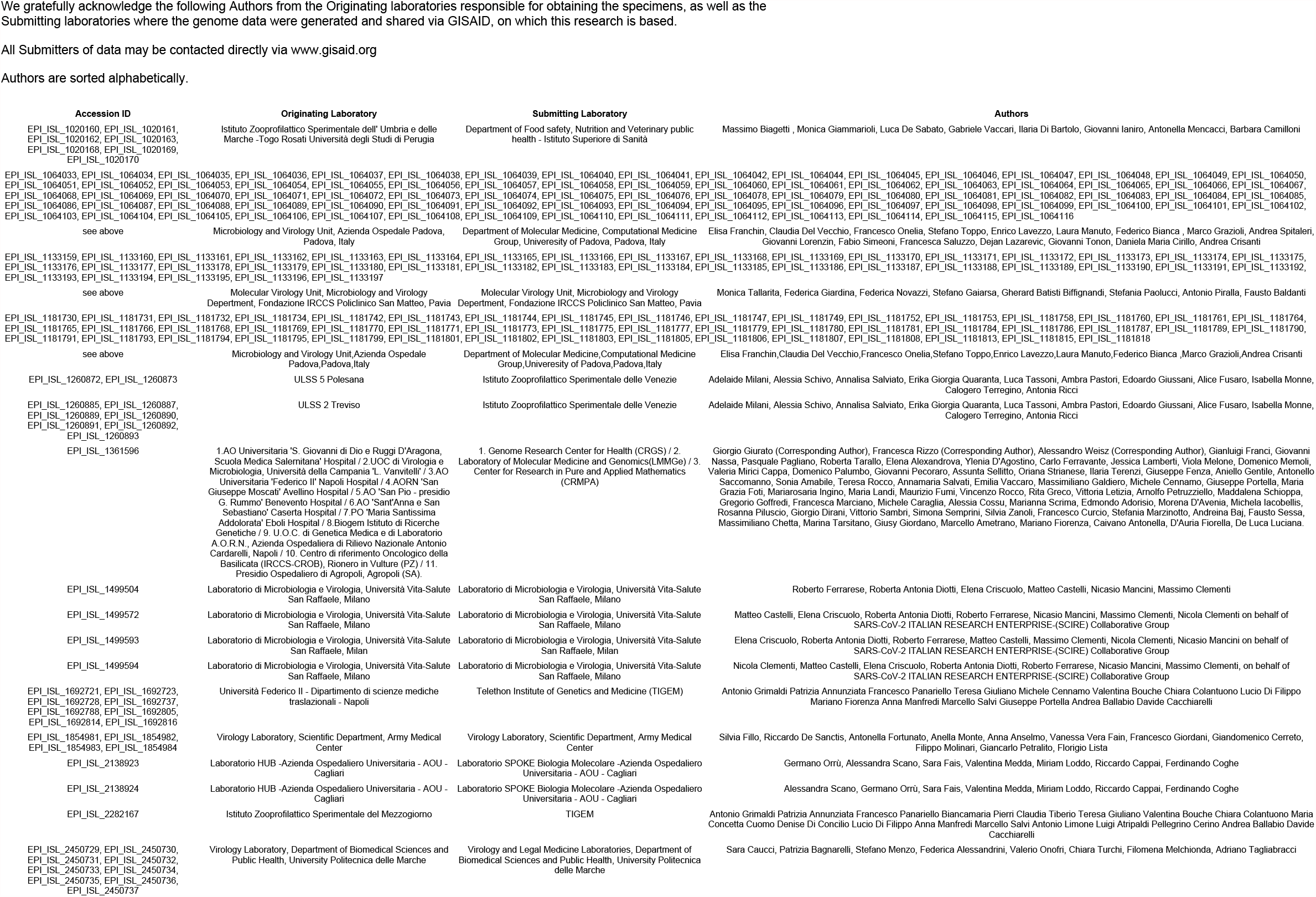
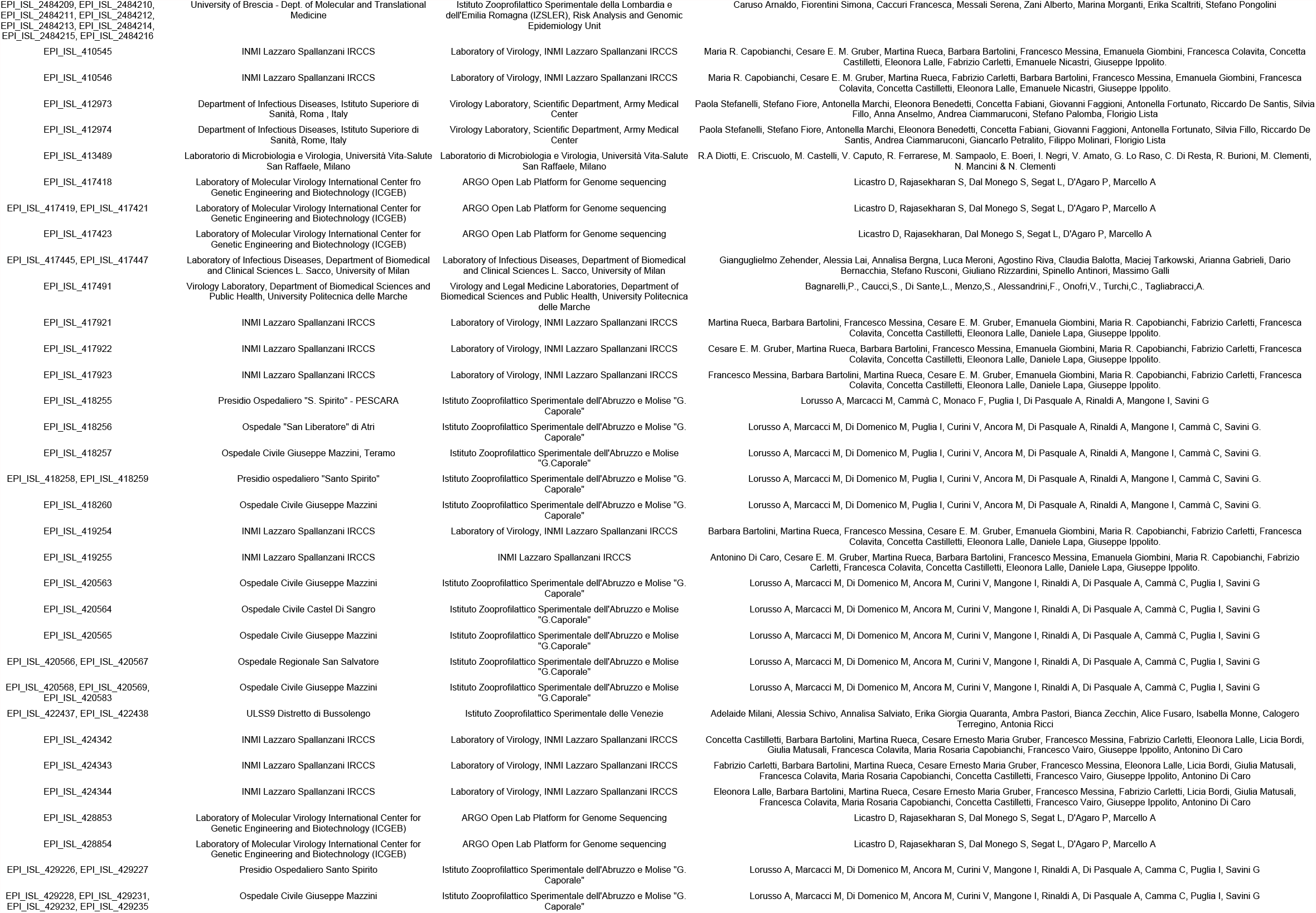
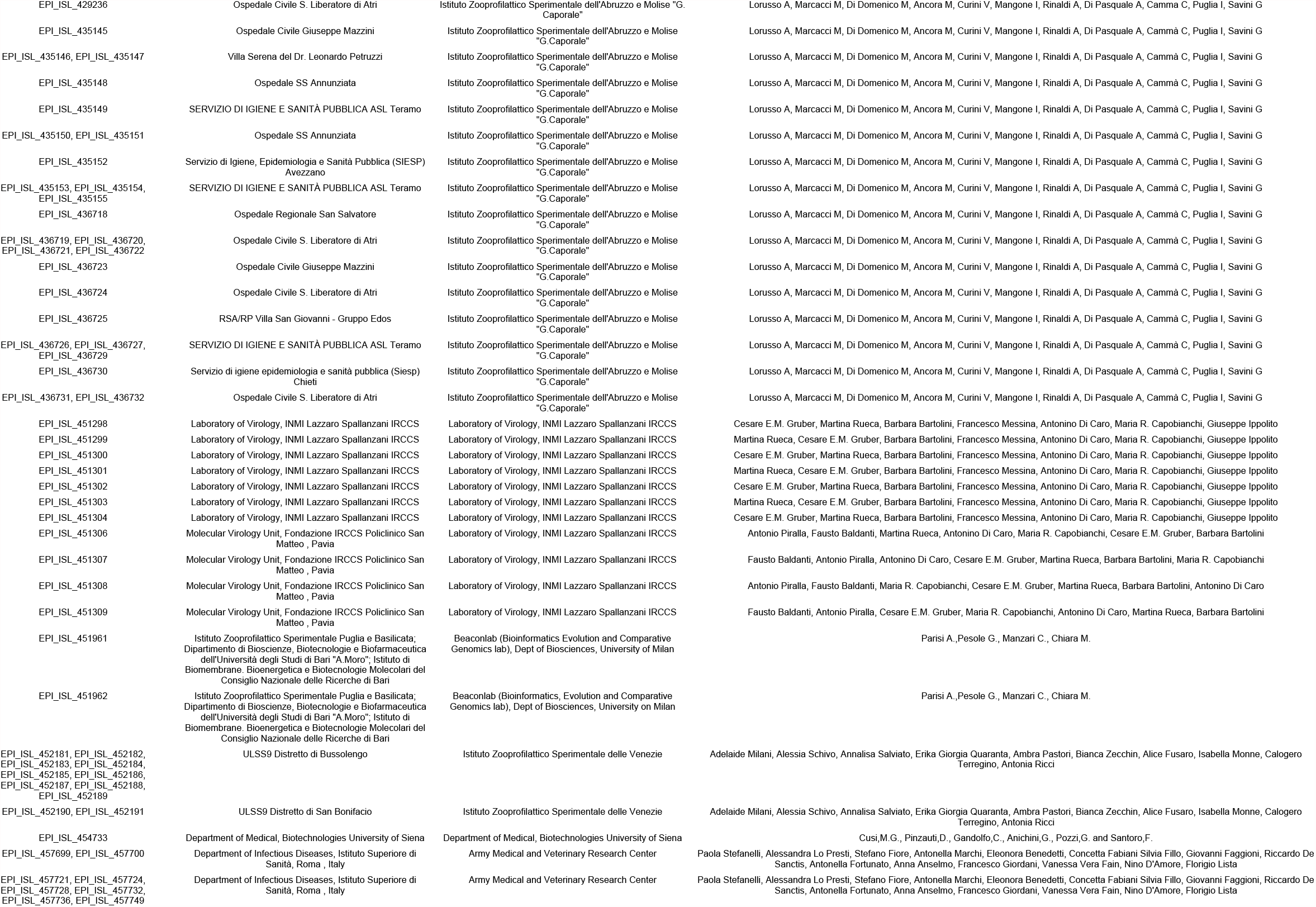
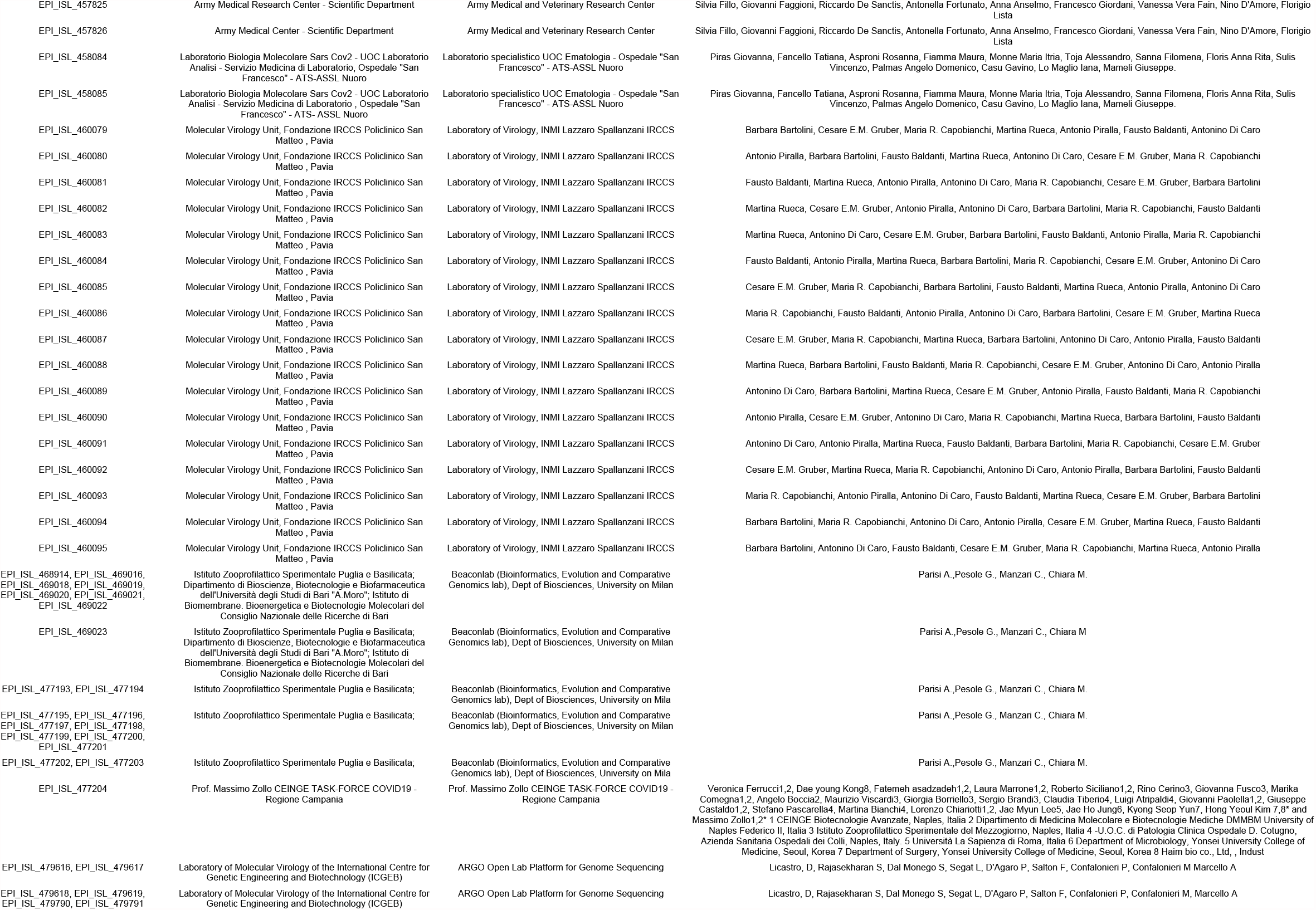
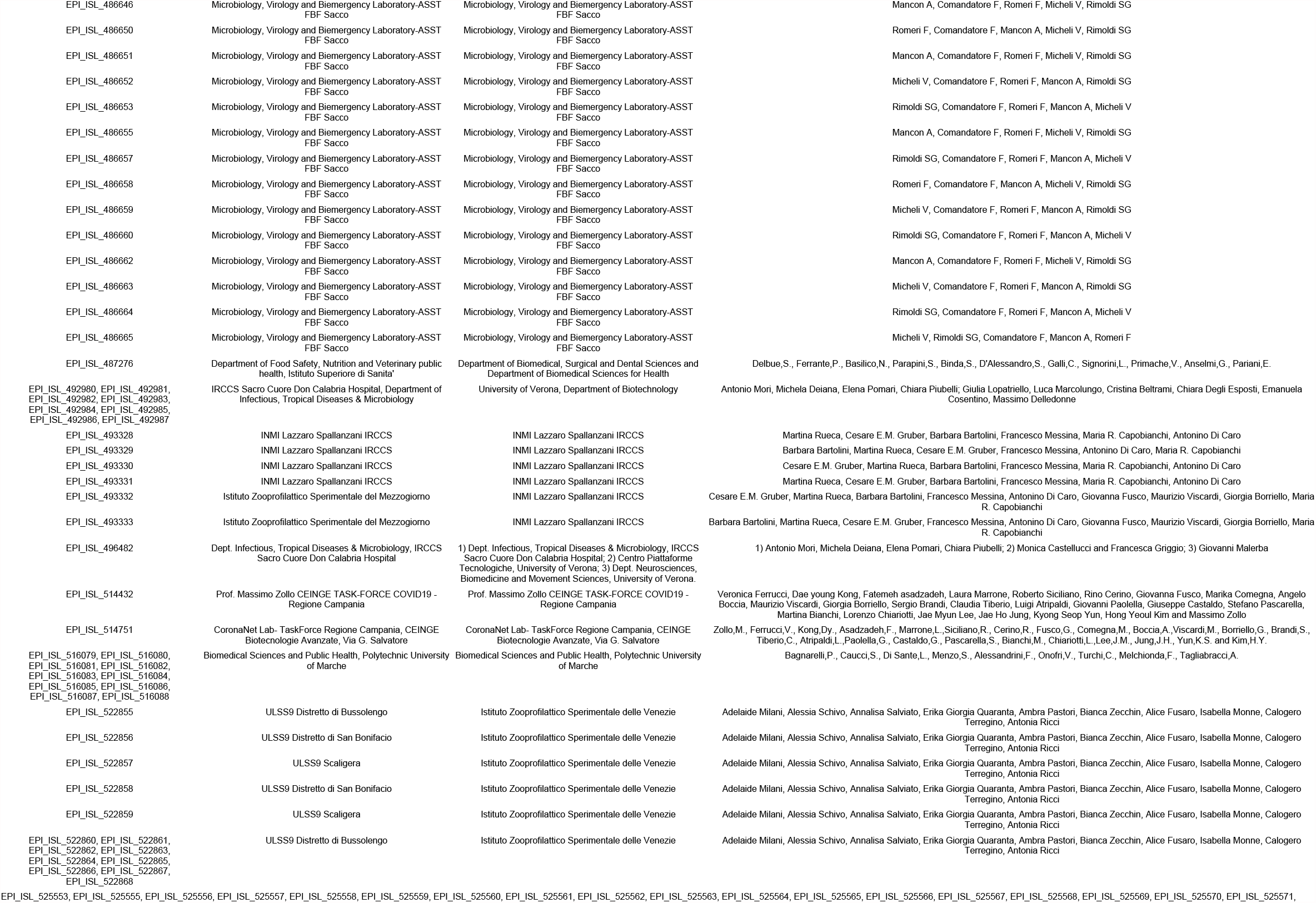
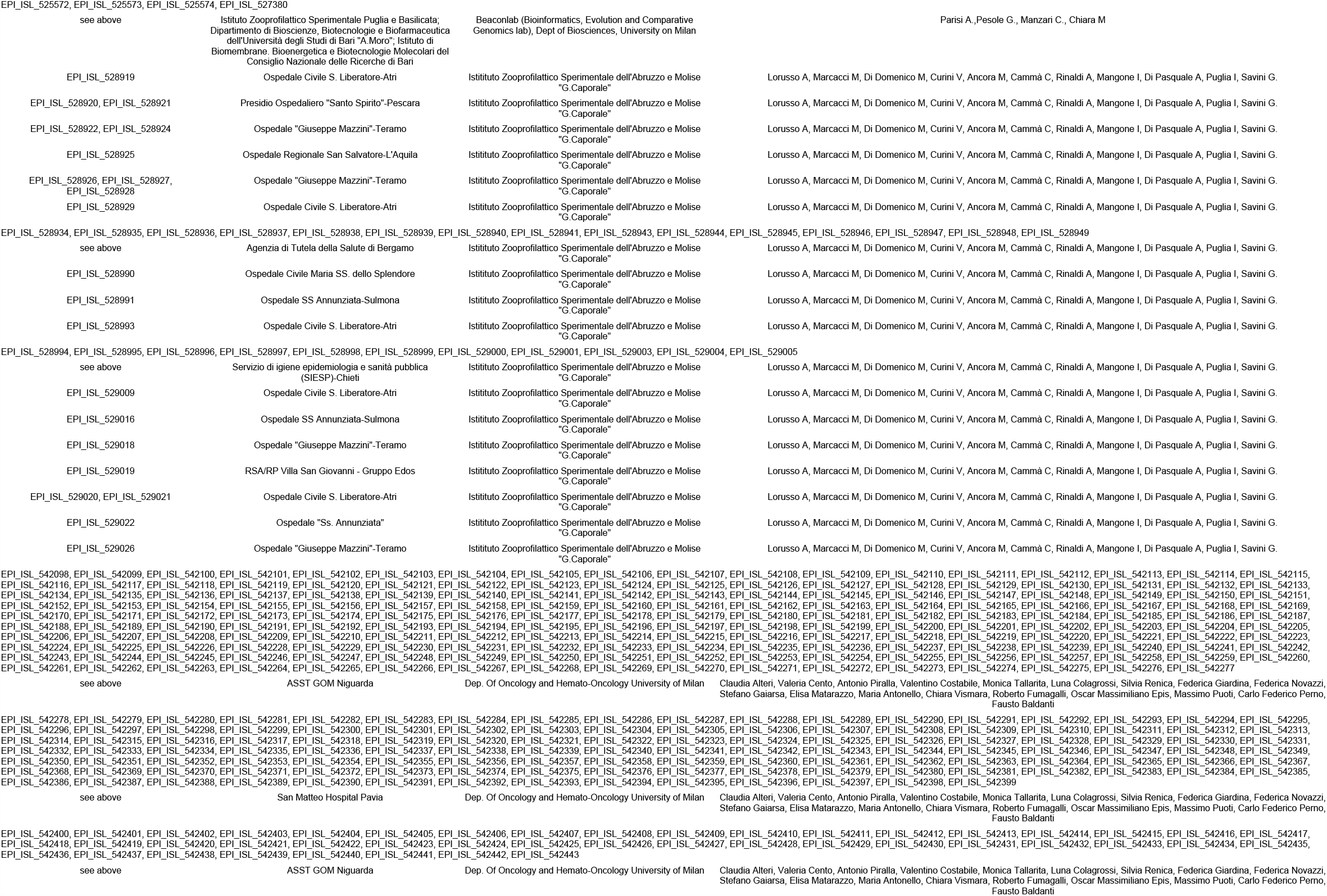
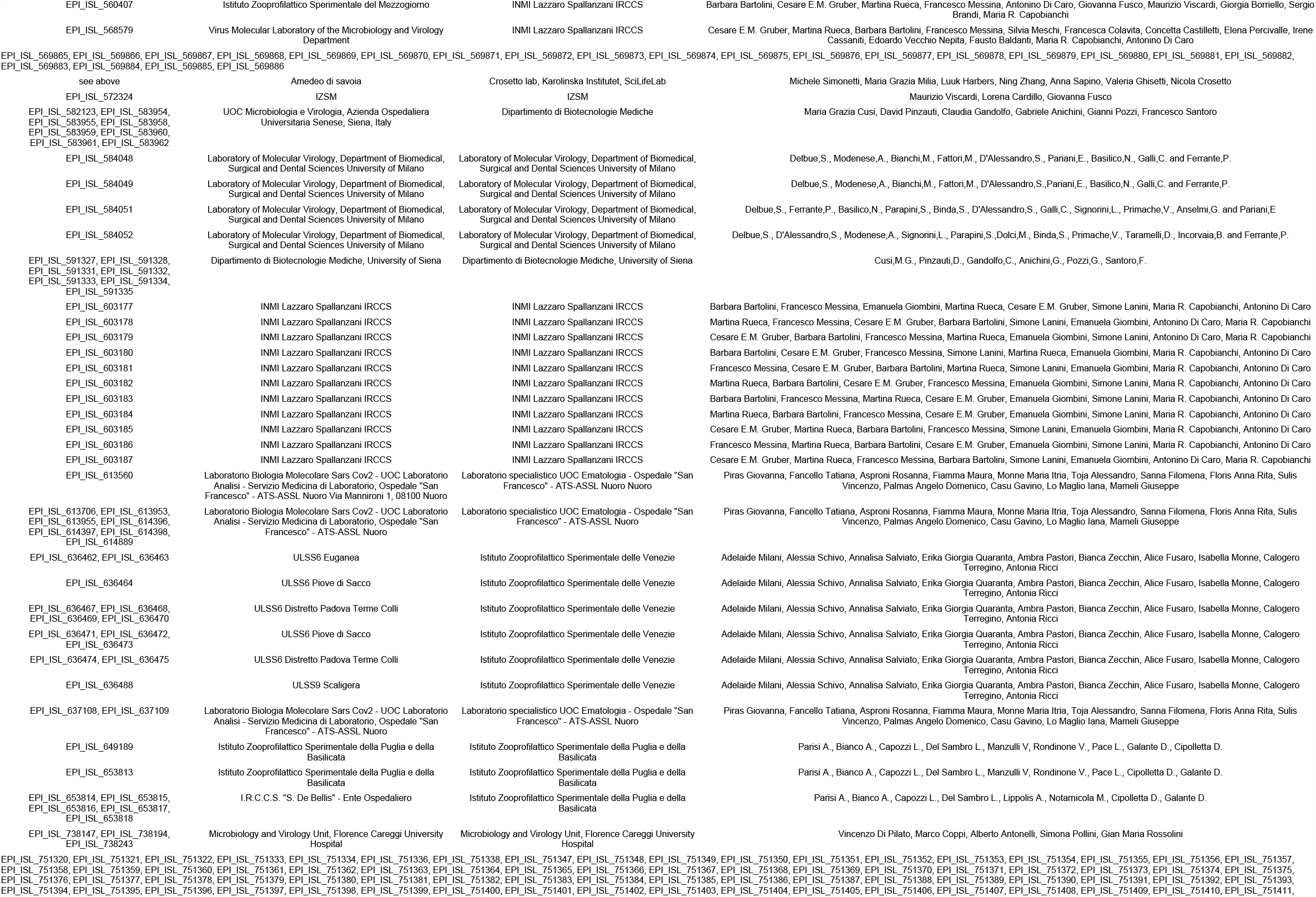
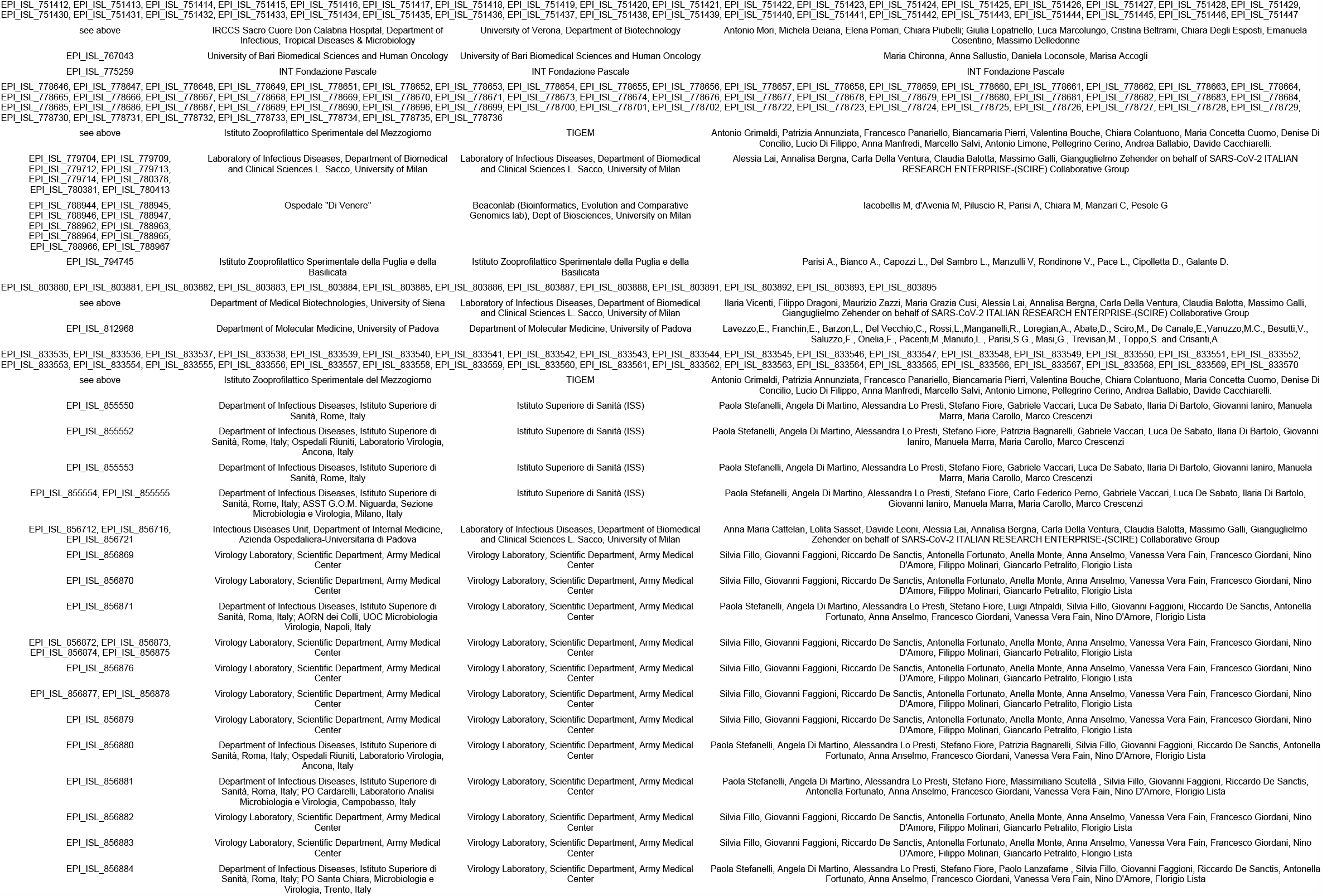
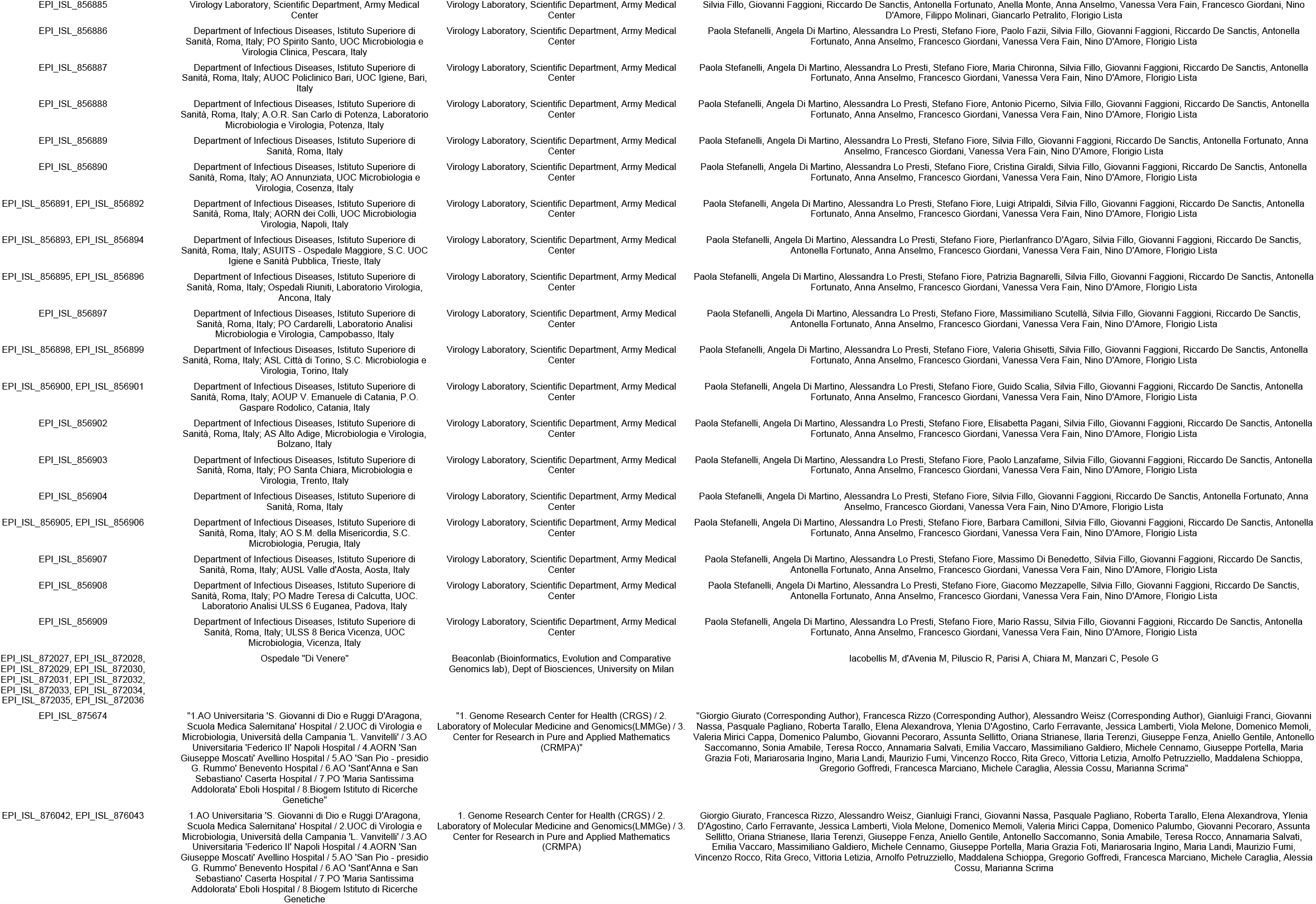
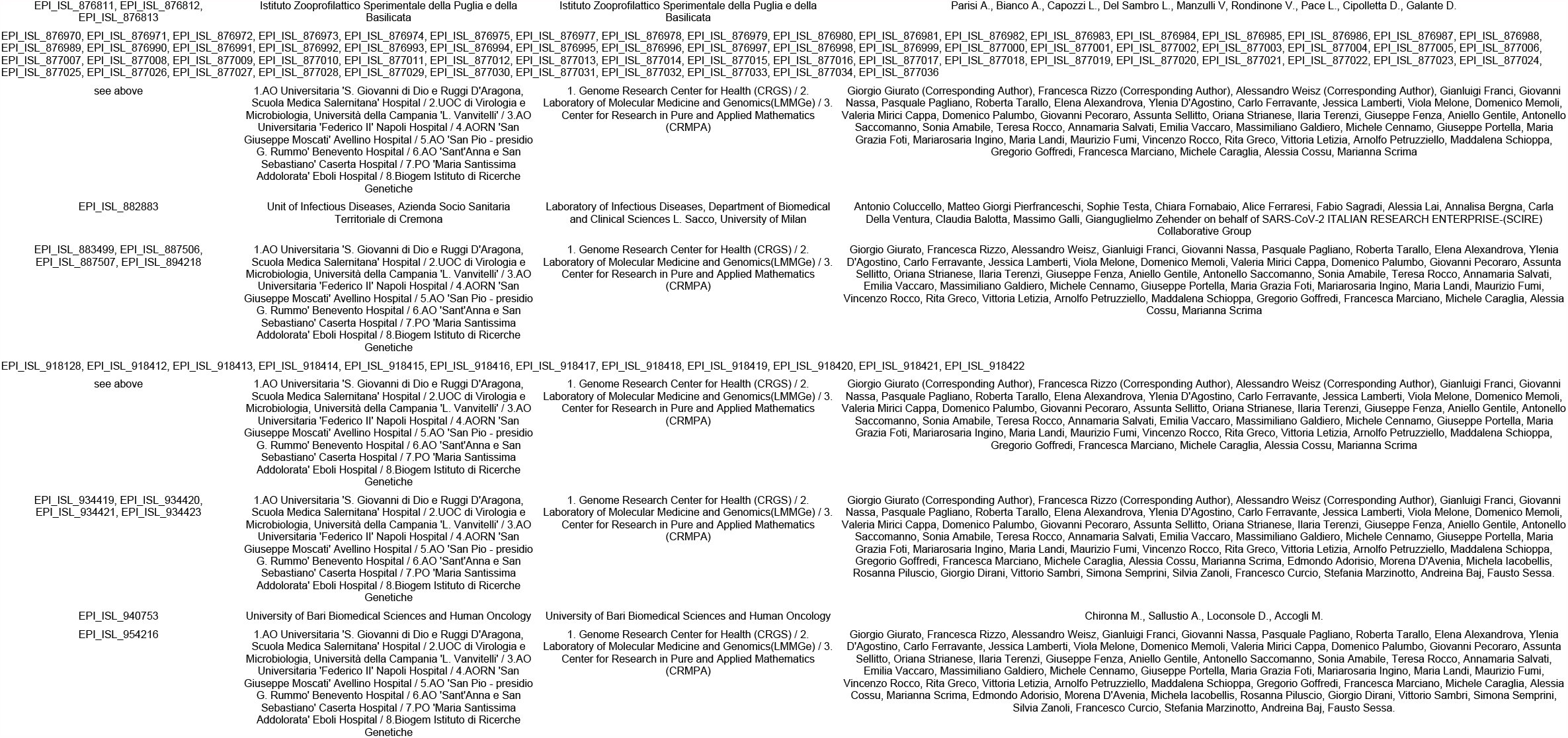

## References

Barido-Sottani, J., T. G. Vaughan, and T. Stadler. 2020. A multitype birth-death model for Bayesian inference of lineage-specific birth and death rates. Systematic Biology 69:973–986.

Beaulieu, J. M. and B.C. O’Meara. 2016. Detecting hidden diversification shifts in models of trait-dependent speciation and extinction. Systematic Biology 65:583–601.

Beerli, P. and J. Felsenstein. 1999. Maximum-likelihood estimation of migration rates and effective population numbers in two populations using a coalescent approach. Genetics 152:763–773.

Boskova, V. and T. Stadler. 2020. PIQMEE: Bayesian phylodynamic method for analysis of large data sets with duplicate sequences. Molecular Biology and Evolution 37:3061–3075.

Cori, A., N. M. Ferguson, C. Fraser, and S. Cauchemez. 2013. A new framework and software to estimate timevarying reproduction numbers during epidemics. American Journal of Epidemiology 178:1505–1512.

Drummond, A. J., G. K. Nicholls, A. G. Rodrigo, and W. Solomon. 2002. Estimating mutation parameters, population history and genealogy simultaneously from temporally spaced sequence data. Genetics 161:1307–1320.

Durrett, R. 2008. Probability models for DNA sequence evolution. Springer.

Etienne, R. S., B. Haegeman, T. Stadler, T. Aze, P. N. Pearson, A. Purvis, and A. B. Phillimore. 2012. Diversity-dependence brings molecular phylogenies closer to agreement with the fossil record. Proceedings of the Royal Society B: Biological Sciences 279:1300–1309.

Ewens, W. J. 1972. The sampling theory of selectively neutral alleles. Theoretical Population Biology 3:87–112.

Ewing, G., G. Nicholls, and A. Rodrigo. 2004. Using temporally spaced sequences to simultaneously estimate migration rates, mutation rate and population sizes in measurably evolving populations. Genetics 168:2407–2420.

Gonzalez-Reiche, A. S., M. M. Hernandez, M. J. Sullivan, B. Ciferri, H. Alshammary, A. Obla, S. Fabre, G. Kleiner, J. Polanco, Z. Khan, et al. 2020. Introductions and early spread of SARS-CoV-2 in the New York city area. Science 369:297–301.

Gupta, A., M. Manceau, T. Vaughan, M. Khammash, and T. Stadler. 2020. The probability distribution of the reconstructed phylogenetic tree with occurrence data. Journal of Theoretical Biology 488:110115.

Hadfield, J., C. Megill, S. M. Bell, J. Huddleston, B. Potter, C. Callender, P. Sagulenko, T. Bedford, and R. A. Neher. 2018. Nextstrain: real-time tracking of pathogen evolution. Bioinformatics 34:4121–4123.

Hein, J., M. Schierup, and C. Wiuf. 2004. Gene genealogies, variation and evolution: a primer in coalescent theory. Oxford university press.

Hermann, W. and J. Leydold. 2014. Generating generalized inverse gaussian random variates. Statistics and Computing 24:547–557.

Hudson, R. R. 1983. Properties of a neutral allele model with intragenic recombination. Theoretical Population Biology 23:183–201.

Karcher, M. D., L. M. Carvalho, M. A. Suchard, G. Dudas, and V. N. Minin. 2020. Estimating effective population size changes from preferentially sampled genetic sequences. PLoS Computational Biology 16:-1007774.

Kendall, D. G. 1948. On the generalized ‘birth-and-death’ process. Ann. Math. Stat. 19:1–15.

Kim, K., R. Omori, and K. Ito. 2017. Inferring epidemiological dynamics of infectious diseases using Tajima’s D statistic on nucleotide sequences of pathogens. Epidemics 21:21–29.

Kingman, J. F. C. 1982. The coalescent. Stochastic processes and their applications 13:235–248.

Kuhner, M. K., J. Yamato, and J. Felsenstein. 1998. Maximum likelihood estimation of population growth rates based on the coalescent. Genetics 149:429–434.

Lanave, C., G. Preparata, C. Sacone, and G. Serio. 1984. A new method for calculating evolutionary substitution rates. Journal of Molecular Evolution 20:86–93.

Lartillot, N. 2006. Conjugate Gibbs sampling for Bayesian phylogenetic models. Journal of Computational Biology 13:1701–1722.

Lartillot, N. and H. Philippe. 2004. A Bayesian mixture model for across-site heterogeneities in the amino-acid replacement process. Molecular Biology and Evolution 21:1095–1109.

Lemey, P., S. Hong, V. Hill, G. Baele, C. Poletto, V. Colizza, A. O’Toole, J. T. McCrone, K. G. Andersen, M. Worobey, M. I. Nelson, A. Rambaut, and M. A. Suchard. 2020. Accommodating individual travel history, global mobility, and unsampled diversity in phylogeography: a SARS-CoV-2 case study. bioRxiv.

Lepage, T., D. Bryant, H. Philippe, and N. Lartillot. 2007. A general comparison of relaxed molecular clock models. Molecular Biology and Evolution 24:2669–2680.

Leventhal, G. E., H. F. Gunthard, S. Bonhoeffer, and T. Stadler. 2013. Using an epidemiological model for phylogenetic inference reveals density dependence in HIV transmission. Molecular Biology and Evolution 31:6–17.

Li, Q., X. Guan, P. Wu, X. Wang, L. Zhou, Y. Tong, R. Ren, K. S. M. Leung, E. H. Y. Lau, J. Y. Wong, et al. 2020. Early transmission dynamics in wuhan, china, of novel coronavirus-infected pneumonia. The New England Journal of Medicine 382:1199–1207.

Maddison, W. P., P. E. Midford, and S. P. Otto. 2007. Estimating a binary character’s effect on speciation and extinction. Systematic Biology 56:701–710.

Maliet, O., F. Hartig, and H. Morion. 2019. A model with many small shifts for estimating species-specific diversification rates. Nature Ecology & Evolution 3:1086–1092.

Manceau, M., A. Gupta, T. Vaughan, and T. Stadler. 2021. The probability distribution of the ancestral population size conditioned on the reconstructed phylogenetic tree with occurrence data. Journal of Theoretical Biology 509:110400.

McDonald, J. H. and M. Kreitman. 1991. Adaptive protein evolution at the Adh locus in Drosophila. Nature 351:652–654.

Morion, H., T. L. Parsons, and J. B. Plotkin. 2011. Reconciling molecular phylogenies with the fossil record. P. Natl. Acad. Sci. USA 108:16327–16332.

Muller, N. F., D. A. Rasmussen, and T. Stadler. 2017. The structured coalescent and its approximations. Molecular Biology and Evolution 34:2970–2981.

Nee, S., R. M. May, and P. H. Harvey. 1994. The reconstructed evolutionary process. Philosophical Transactions of the Royal Society of London B: Biological Sciences 344:305–311.

Novembre, J., T. Johnson, K. Bryc, Z. Kutalik, A. R. Boyko, A. Auton, A. Indap, K. S. King, S. Bergmann, M. R. Nelson, et al. 2008. Genes mirror geography within Europe. Nature 456:98–101.

Parag, K. V., L. du Plessis, and O. G. Pybus. 2020. Jointly inferring the dynamics of population size and sampling intensity from molecular sequences. Molecular Biology and Evolution 37:2414–2429.

Parag, K. V., O. G. Pybus, and C.-H. Wu. 2021. Are skyline plot-based demographic estimates overly dependent on smoothing prior assumptions? Systematic Biology.

Pybus, O. G., A. J. Drummond, T. Nakano, B. H. Robertson, and A. Rambaut. 2003. The epidemiology and iatrogenic transmission of hepatitis C virus in Egypt: a Bayesian coalescent approach. Molecular Biology and Evolution 20:381–387.

Pybus, O. G. and P. H. Harvey. 2000. Testing macro-evolutionary models using incomplete molecular phylogenies. P. Roy. Soc. Lend. B. Bio. 267:2267–2272.

Rasmussen, D. A., O. Ratmann, and K. Koelle. 2011. Inference for nonlinear epidemiological models using genealogies and time series. PLoS Computational Biology 7:-1002136.

Skoglund, P., P. Sjodin, T. Skoglund, M. Lascoux, and M. Jakobsson. 2014. Investigating population history using temporal genetic differentiation. Molecular Biology and Evolution 31:2516–2527.

Stadler, T. 2010. Sampling-through-time in birth-death trees. Journal of Theoretical Biology 267:396–404.

Stadler, T. 2011. Mammalian phylogeny reveals recent diversification rate shifts. P. Natl. Acad. Sci. USA 108:61876192.

Stadler, T., D. Kiihnert, S. Bonhoeffer, and A. J. Drummond. 2013. Birth-death skyline plot reveals temporal changes of epidemic spread in HIV and hepatitis C virus (HCV). Proceedings of the National Academy of Sciences 110:228–233.

Stephens, M. and P. Donnelly. 2000. Inference in molecular population genetics. Journal of the Royal Statistical Society: Series B (Statistical Methodology) 62:605–635.

Taits, S., M. Betancourt, D. Simpson, A. Vehtari, and A. Gelman. 2018. Validating Bayesian inference algorithms with simulation-based calibration. arXiv:1804.06788 [stat] ArXiv: 1804.06788.

Tavare, S. 2004. Part I: Ancestral inference in population genetics. Pages 1–188 in Lectures on probability theory and statistics. Springer.

Vaughan, T. G., D. Kuhnert, A. Popinga, D. Welch, and A. J. Drummond. 2014. Efficient Bayesian inference under the structured coalescent. Bioinformatics 30:2272–2279.

Vaughan, T. G., G. E. Leventhal, D. A. Rasmussen, A. J. Drummond, D. Welch, and T. Stadler. 2019. Estimating epidemic incidence and prevalence from genomic data. Molecular Biology and Evolution 36:1804–1816.

Vaughan, T. G., J. Scire, S. A. Nadeau, and T. Stadler. 2020. Estimates of outbreak-specific SARS-CoV-2 epidemiological parameters from genomic data. medRxiv.

Volz, E. M., S. L. K. Pond, M. J. Ward, A. J. L. Brown, and S. D. Frost. 2009. Phylodynamics of infectious disease epidemics. Genetics 183:1421–1430.

